# A single-dose live-attenuated YF17D-vectored SARS-CoV2 vaccine candidate

**DOI:** 10.1101/2020.07.08.193045

**Authors:** Lorena Sanchez Felipe, Thomas Vercruysse, Sapna Sharma, Ji Ma, Viktor Lemmens, Dominique van Looveren, Mahadesh Prasad Arkalagud Javarappa, Robbert Boudewijns, Bert Malengier-Devlies, Suzanne F. Kaptein, Laurens Liesenborghs, Carolien De Keyzer, Lindsey Bervoets, Madina Rasulova, Laura Seldeslachts, Sander Jansen, Michael Bright Yakass, Osbourne Quaye, Li-Hsin Li, Xin Zhang, Sebastiaan ter Horst, Niraj Mishra, Lotte Coelmont, Christopher Cawthorne, Koen Van Laere, Ghislain Opdenakker, Greetje Van de Velde, Birgit Weynand, Dirk E. Teuwen, Patrick Matthys, Johan Neyts, Hendrik Jan Thibaut, Kai Dallmeier

## Abstract

The explosively expanding COVID-19 pandemic urges the development of safe, efficacious and fast-acting vaccines to quench the unrestrained spread of SARS-CoV-2. Several promising vaccine platforms, developed in recent years, are leveraged for a rapid emergency response to COVID-19^1^. We employed the live-attenuated yellow fever 17D (YF17D) vaccine as a vector to express the prefusion form of the SARS-CoV-2 Spike antigen. In mice, the vaccine candidate, tentatively named YF-S0, induces high levels of SARS-CoV-2 neutralizing antibodies and a favorable Th1 cell-mediated immune response. In a stringent hamster SARS-CoV-2 challenge model^2^, vaccine candidate YF-S0 prevents infection with SARS-CoV-2. Moreover, a single dose confers protection from lung disease in most vaccinated animals even within 10 days. These results warrant further development of YF-S0 as a potent SARS-CoV-2 vaccine candidate.

## Vaccine design and rationale

Protective immunity against SARS-CoV-2 and other coronaviruses is believed to depend on neutralizing antibodies (nAbs) targeting the viral Spike (S) protein ^3,4^. In particular, nAbs specific for the N-terminal S1 domain containing the Angiotensin Converting Enzyme 2 (ACE2) receptor binding domain (RBD) interfere with and have been shown to prevent viral infection in several animal models^5,6^.

The live-attenuated YF17D vaccine is known for its outstanding potency to rapidly induce broad multi-functional innate, humoral and cell-mediated immunity (CMI) responses that may result in life-long protection following a single vaccine dose in nearly all vaccinees ^7,8^. These favorable characteristics of the YF17D vaccine translate also to vectored vaccines based on the YF17D backbone ^9^. YF17D is used as viral vector in two licensed human vaccines [Imojev^®^ against Japanese encephalitis virus (JEV) and Dengvaxia^®^ against dengue virus (DENV)]. For these two vaccines, genes encoding the YF17D surface antigens prM/E, have been swapped with those of JEV or DENV, respectively. Potent Zika virus vaccines based on this ChimeriVax approach are in preclinical development ^10^.

YF17D is a small (+)-ss RNA live-attenuated virus with a limited vector capacity, but it has been shown to tolerate insertion of foreign antigens at two main sites in the viral polyprotein ^11^. Importantly, an insertion of foreign sequences is constrained by (*i*) the complex topology and post-translational processing of the YF17D polyprotein; and, (*ii*) the need to express the antigen of interest in an immunogenic, likely native, fold, to yield a potent recombinant vaccine.

Using an advanced reverse genetics system for the generation of recombinant flaviviruses^12,13^, a panel of YF17D-based COVID-19 vaccine candidates (YF-S) was designed. These express codon-optimized versions of the S protein [either in its native cleavable S1/2, or non-cleavable prefusion S0 conformation or its S1 subdomain] of the prototypic SARS-CoV-2 Wuhan-Hu-1 strain (GenBank: MN908947.3), as in-frame fusion within the YF17D-E/NS1 intergenic region (YF-S1/2, YF-S0 and YF-S1) (Figure 1A, Figure S1). As outlined below, variant YF-S0 was finally selected as lead vaccine candidate based on its superior immunogenicity, efficacy and favorable safety profile.

**Fig. 1.**
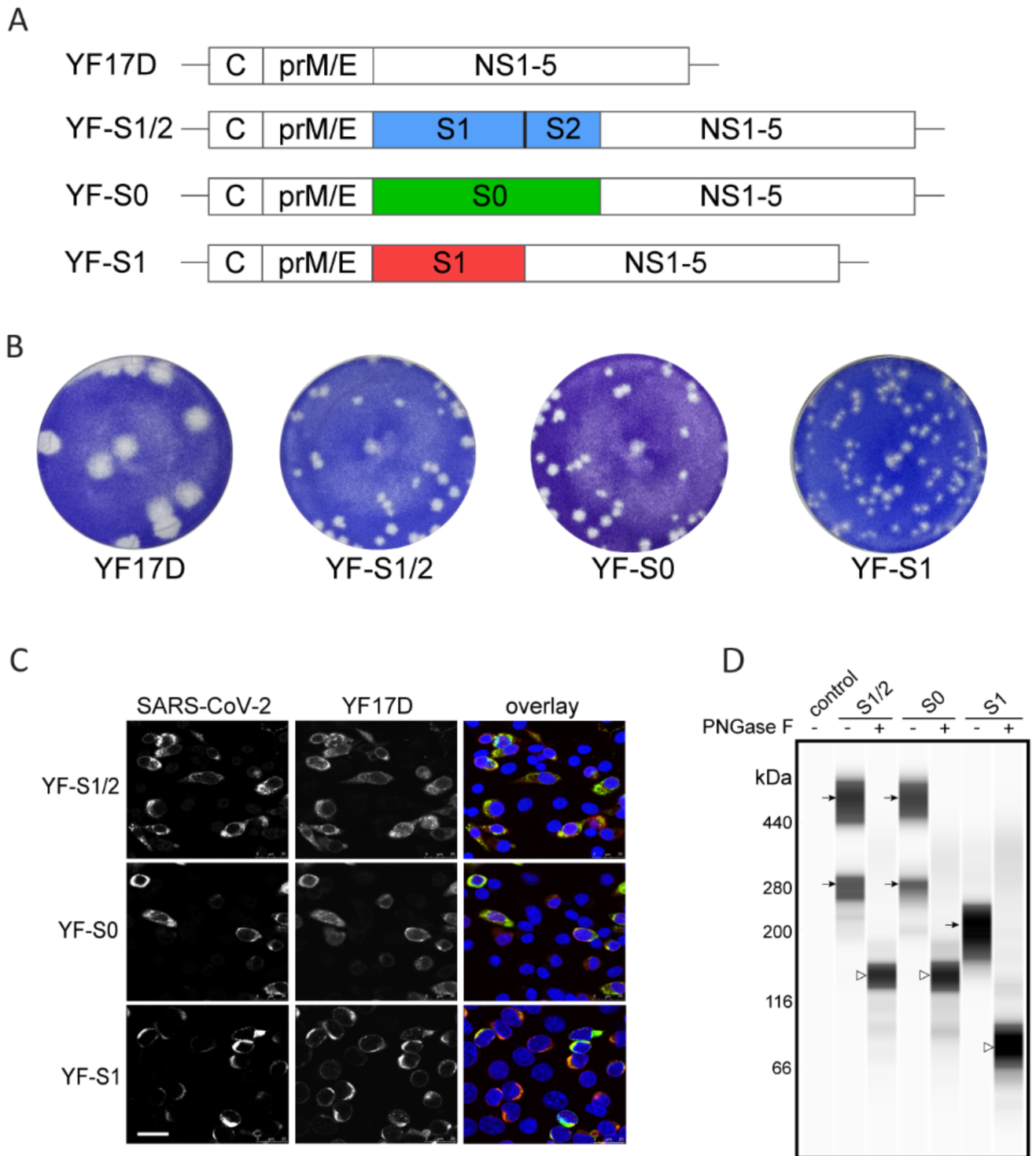
Vaccine design and antigenicity. **(A)** Schematic representation of YF17D-based SARS-CoV-2 vaccine candidates (YF-S). YF-S1/2 expresses the native cleavable post-fusion form of the S protein (S1/2), YF-S0 the non-cleavable pre-fusion conformation (S0), and YF-S1 the N-terminal (receptor binding domain) containing S1 subunit of the S protein. For molecular details in vaccine design see Methods section. **(B)** Representative pictures of plaque phenotypes from different YF-S vaccine constructs on BHK-21 cells in comparison to YF17D. **(C)** Confocal immunofluorescent images of BHK-21 cells three days post-infection with different YF-S vaccine constructs staining for SARS-CoV-2 Spike antigen (green) and YF17D (red) (nuclei stained with DAPI, blue; white scalebar: 25 µm). **(D)** Immunoblot analysis of SARS-CoV-2 Spike (S1/2, S0 and S1) antigen and SARS Spike expression after transduction of BHK-21 cells with different YF-S vaccine candidates. Prior to analysis, cell lysates were treated with PNGase F to remove their glycosylation or left untreated (black arrows – glycosylated forms of S; white arrows – de-glycosylated forms).

Infectious live-attenuated YF-S viruses were rescued by plasmid transfection into baby hamster kidney (BHK-21) cells. Transfected cells presented with a virus-induced cytopathic effect; infectious virus progeny was recovered from the supernatant and further characterized. Each construct results in a unique plaque phenotype, smaller than that of the parental YF17D (Figure 1B), in line with some replicative trade-off posed by the inserted foreign sequences. S or S1 as well as YF17D antigens were readily visualized by double staining of YF-S infected cells with SARS-CoV-2 Spike and YF17D-specific antibodies (Figure 1C). The expression of S or S1 by the panel of YF-S variants was confirmed by immunoblotting of lysates of freshly infected cells. Treatment with PNGase F allowed to demonstrate a proper glycosylation pattern (Figure 1D).

In line with a smaller plaque phenotype, intracranial (i.c.) inoculation of YF17D or the YF-S variants in suckling mice confirmed the attenuation of the different YF-S as compared to the empty vector YF17D (Figure 2A and B and Figure S2). Mouse pups inoculated i.c. with 100 plaque forming units (PFU) of the parental YF17D stopped growing (Figure S2A) and succumbed to infection within seven days (median day of euthanasia; MDE) (Figure 2A), whereas pups inoculated with the YF-S variants continued to grow. From the group inoculated with YF-S0 only half needed to be euthanized (MDE 17,5 days). For the YF-S1/2 and YF-S1 groups MDE was 12 and 10 days respectively; thus in particular YF-S0 has a markedly reduced neurovirulence. Likewise, YF-S0 is also highly attenuated in type I and II interferon receptor deficient AG129 mice, that are highly susceptible to (a neurotropic) YF17D infection^13,14^.

**Fig. 2.**
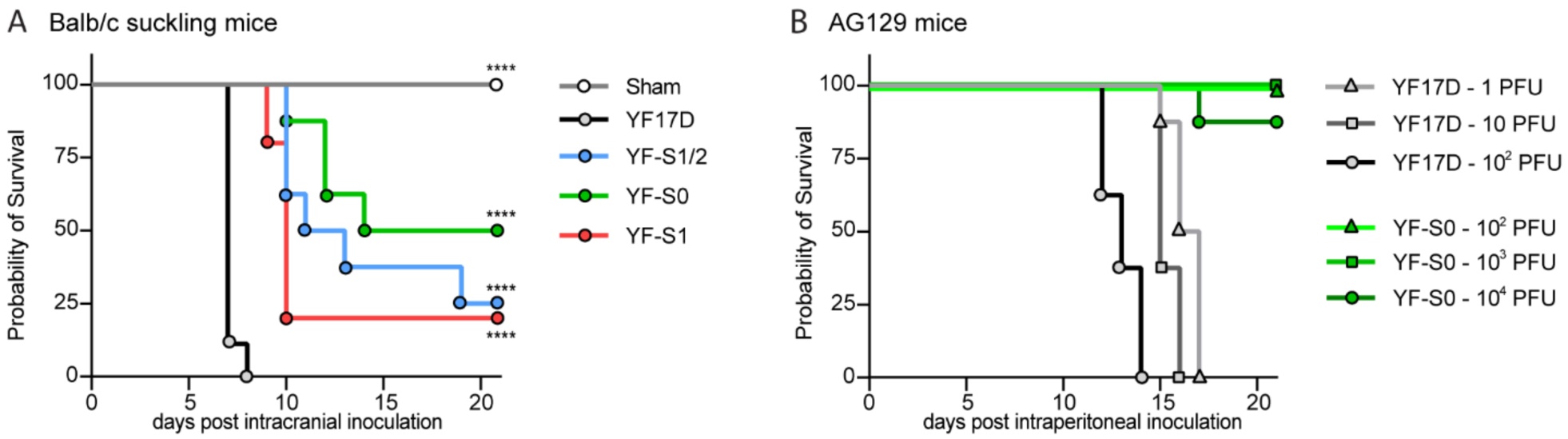
Attenuation of YF-S vaccine candidates. **(A)** Survival curve of suckling Balb/c mice (up to 21 days) after intracranial (i.c.) inoculation with 100 PFU of vaccine candidates YF-S1/2 (n=8, blue), YF-S0 (n=8, green), YF-S1 (n=8, red) in comparison to sham (n=10, grey) or YF17D (n=9, black). **(B)** Survival curve of AG129 mice (up to 21 days) after intraperitoneal (i.p.) inoculation with a dose range of YF-S0 (10^2^, 10^3^ and 10^4^ PFU; green) in comparison to YF17D (1, 10 and 10^2^ PFU black and grey). Statistical significance between groups was calculated by the Log-rank Mantel-Cox test (**** P < 0.0001).

Whereas 1 PFU of YF17D resulted in neuro-invasion requiring euthanasia of all mice (MDE 16 days) (Figure 2B), a 1000-fold higher inoculum of YF-S0 did not result in any disease (Figure S2C) and only 1 in 12 animals that received a 10,000 higher inoculum needed to be euthanized (Figure 2B). In summary, a set of transgenic replication-competent YF17D variants (YF-S) was generated that express different forms of the SARS-CoV-2 S antigen and that are highly attenuated in mice in terms of neurovirulence and neurotropism as compared to YF17D.

### Immunogenicity and protection against SARS-CoV-2 infection and COVID-19-like disease in a stringent hamster model

To assess the potency of the various vaccine constructs, a stringent hamster challenge model was developed^2^. Animals were vaccinated at day 0 with 10^3^ PFU (i.p. route) of the different constructs or the negative controls and were boosted 7 days later with the same dose (Figure 3A). At day 21 post-vaccination, all hamsters vaccinated with YF-S1/2 and YF-S0 (n=12 from two independent experiments) had seroconverted to high levels of S-specific IgG and virus nAbs (Figure 3B,C; see Figure S3 for benchmarking of SARS-CoV-2 serum neutralization test, SNT). For YF-S1/2 log_10_ geometric mean titers (GMT) for IgG and nAbs were 3.2 (95% CI, 2.9-3.5) and 1.4 (95% CI, 1.1-1.9) respectively, while in the case of YF-S0 GMT values for IgG and nAbs of 3.5 (95% CI, 3.3-3.8) and 2.2 (95% CI, 1.9-2.6) were measured, with rapid seroconversion kinetics (50% seroconversion rate < 2 weeks; Figure 3D). By contrast, only 1 out of 12 hamsters that had received YF-S1 seroconverted and this with a low level of nAbs. This indicates the need for a full-length S antigen to elicit an adequate humoral immune response.

**Fig. 3.**
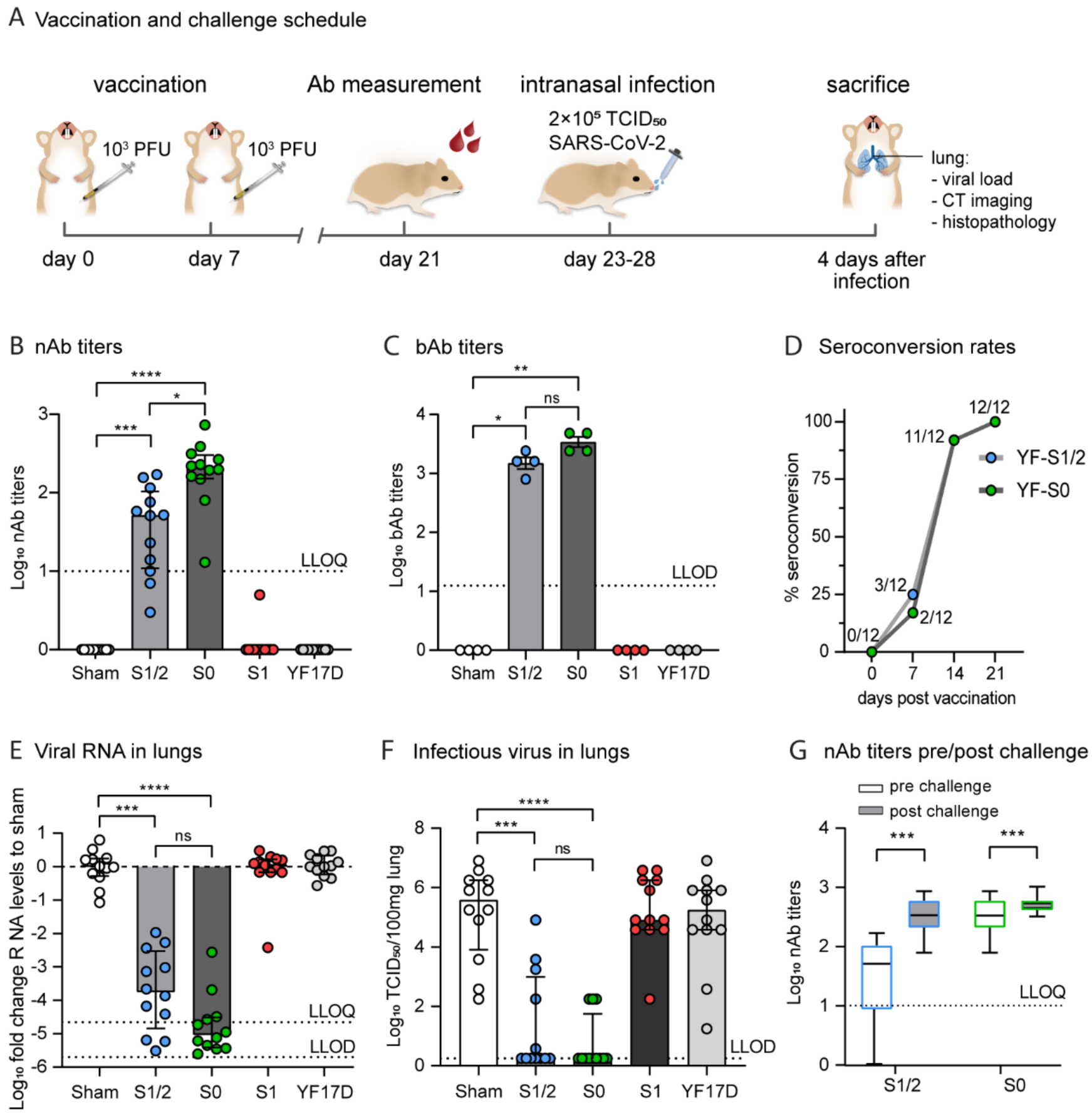
Immunogenicity and protective efficacy of YF-S vaccine candidates in hamsters. **(A) Schematic representation of vaccination and challenge schedule**. Syrian hamsters were immunized twice i.p. at day 0 and 7 with 10^3^ PFU each of vaccine constructs YF-S1/2 (blue, n=12), YF-S0 (green, n=12), YF-S1 (red, n=12), sham (white, n=12) or YF17D (grey, n=12) (two independent experiments). Subsequently, animals were intranasally inoculated with 2 × 10^5^ TCID_50_ of SARS-CoV-2 and followed up for four days. **(B-D) Humoral immune responses**. Neutralizing antibodies (nAb) **(B)** and total binding IgG (bAb) **(C)** in hamsters vaccinated with different vaccine candidates (sera collected at day 21 post-vaccination in both experiments; minipools of sera of three animals each were analyzed for quantification of bAb). **(D)** Seroconversion rates at indicated days post-vaccination with YF-S1/2 and YF-S0 (number of animals with detectable bAbs at each time point are referenced). **(E, F) Protection from SARS-CoV-2 infection**. Viral loads in lungs of hamsters four days after intranasal infection were quantified by RT-qPCR **(E)** and virus titration **(F)**. Viral RNA levels were determined in the lungs, normalized against β-actin and fold-changes were calculated using the 2^(-ΔΔCq)^ method compared to the median of sham-vaccinated hamsters. Infectious viral loads in the lungs are expressed as number of infectious virus particles per 100 mg of lung tissue. **(G) Anamnestic response**. Comparison of the levels of nAbs prior and four days after challenge. For a pairwise comparison of responses in individual animals see Supplementary Figure S4C and D. Dotted line indicating lower limit of quantification (LLOQ) or lower limit of detection (LLOD) as indicated. Data shown are medians ± IQR. Statistical significance between groups was calculated by the non-parametric ANOVA, Kruskall-Wallis with uncorrected Dunn’s test (B-F), or a non-parametric two-tailed Wilcoxon matched-pairs rank test (G) (ns = Non-Significant, P > 0.05, * P < 0.05, ** P < 0.01, *** P < 0.001, **** P<0.0001).

Next, vaccinated hamsters were challenged intranasally (either at day 23 or day 28 post vaccination) with 2 × 10^5^ PFU of SARS-CoV-2. At day 4 post-infection, high viral loads were detected in lungs of sham-vaccinated controls and animals vaccinated with YF17D as matched placebo (Figure 4A, B). Infection was characterized by a severe lung pathology with multifocal necrotizing bronchiolitis, leukocyte infiltration and edema, resembling findings in patients with severe COVID-19 bronchopneumonia (Figure 4A specimen pictures and 4B radar plot; Supplementary-histogram). By contrast, hamsters vaccinated with YF-S0 were protected against this aggressive challenge (Figure 3E-F). As compared to sham-vaccinated controls, YF-S0 vaccinated animals had a median reduction of 5 log_10_ (IQR, 4.5-5.4) in viral RNA loads (p <0.0001; Figure 3D), and of 5.3 log_10_ (IQR, 3.9-6.3) for infectious SARS-CoV-2 virus in the lungs (p <0.0001; Figure 3E). Moreover, infectious virus was no longer detectable in 10 of 12 hamsters (two independent experiments), and viral RNA was reduced to non-quantifiable levels in their lungs. Residual RNA measured in 2 out of 12 animals may equally well represent residues of the high-titer inoculum as observed in non-human primate models^15-18^. Vaccination with YF-S0 (two doses of 10^3^ PFU) also efficiently prevented systemic viral dissemination; in most animals, no or only very low levels of viral RNA were detectable in spleen, liver, kidney and heart four days after infection (Figure S4A). Similarly and in full support, a slightly different dose and schedule used for vaccination (5 × 10^3^ PFU of YF-S0 at day 0 and 7 respectively) resulted in all vaccinated hamsters (n=7) in respectively a 6 log_10_ (IQR, 4.6-6.6) and 5.7 log_10_ (IQR, 5.7-6.6) reduction of viral RNA and infectious virus titers as compared to sham (Figure S5). Finally, vaccination with YF-S0 may induce saturating levels of nAbs thereby conferring sterilizing immunity, as demonstrated by the fact that in about half of the YF-S0 vaccinated hamsters no anamnestic antibody response was observed following challenge (Figure 3G and S4B-D (paired nAb analysis)). By contrast, in hamsters vaccinated with the second-best vaccine candidate YF-S1/2, nAb levels further increased following SARS-CoV-2 infection (in 11 out of 12 animals) whereby a plateau was only approached after challenge.

**Fig. 4.**
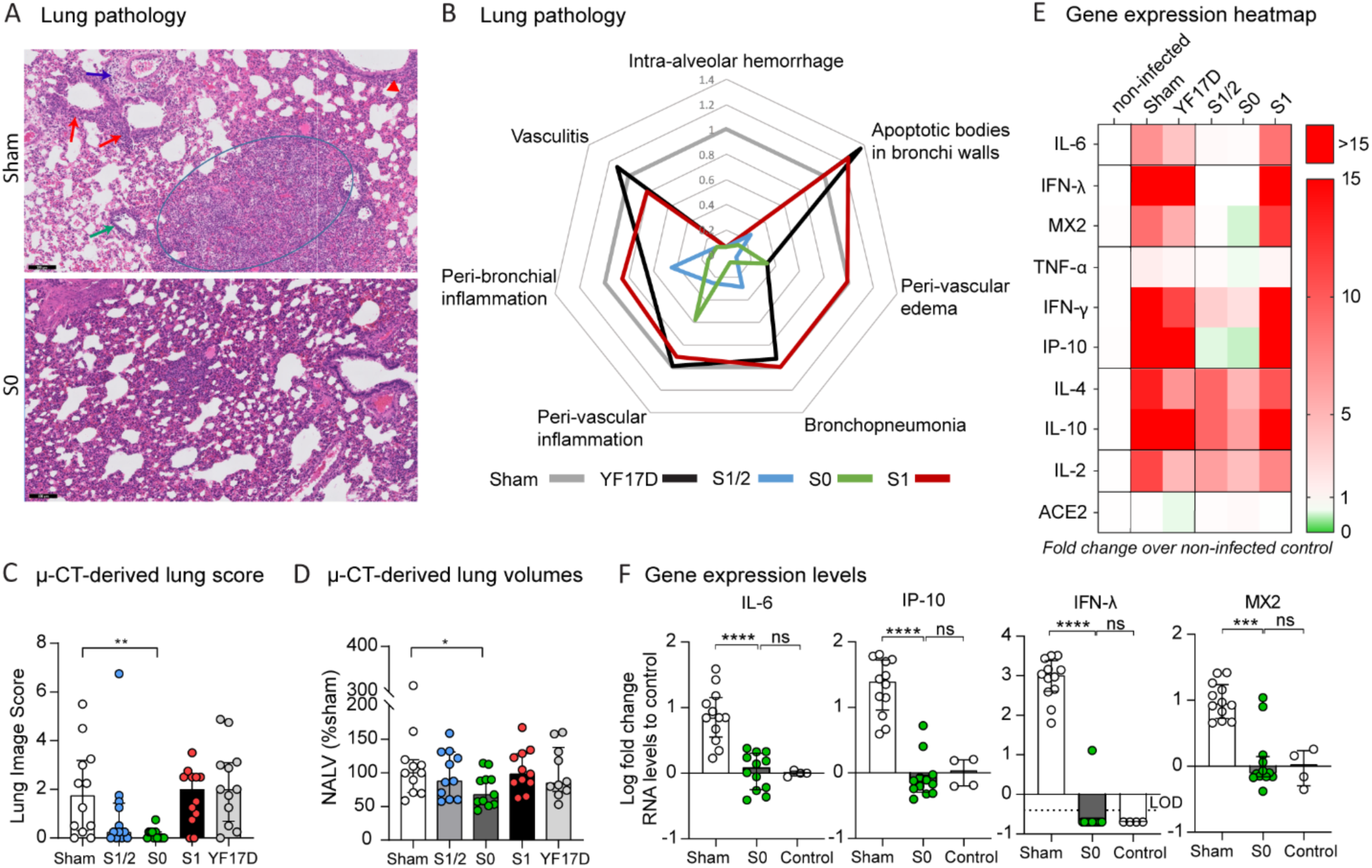
Protection from lung disease in YF-S vaccinated hamsters. **(A) Representative H&E images** of the lungs of a diseased (sham-vaccinated and infected) and a YF-S0-vaccinated and challenged hamster. Peri-vascular edema (blue arrow); peri-bronchial inflammation (red arrows); peri-vascular inflammation (green arrow); bronchopneumonia (circle), apoptotic body in bronchial wall (red arrowhead). **(B)** A spider-web plot showing histopathological score for signs of lung damage (peri-vascular edema, bronchopneumonia, peri-vascular inflammation, peri-bronchial inflammation, vasculitis, intra-alveolar hemorrhage and apoptotic bodies in bronchus walls) normalized to sham (grey). Black scalebar: 100 µm **(C-D) Micro-CT-derived analysis of lung disease**. Five transverse cross sections at different positions in the lung were selected for each animal and scored to quantify lung consolidations **(C)** or used to quantify the non-aerated lung volume (NALV) **(D)**, as functional biomarker reflecting lung consolidation. **(E) Heat-map** showing differential expression of selected antiviral, pro-inflammatory and cytokine genes in lungs of sham- or YF-S-vaccinated hamsters after SARS-CoV-2 challenge four days p.i. (n=12 per treatment group) relative to non-treated non-infected controls (n=4) (scale represents fold-change over controls). RNA levels were determined by RT-qPCR on lung extracts, normalized for β-actin mRNA levels and fold-changes over the median of uninfected controls were calculated using the 2^(-ΔΔCq)^ method. **(F) Individual expression profiles of mRNA levels** of IL-6, IP-10, IFN-λ and MX2, with data presented as median ± IQR relative to the median of non-treated non-infected controls. For IFN-λ, where all control animals had undetectable RNA levels, fold-changes were calculated over the lowest detectable value (LLOD – lower limit of detection; dotted line). Statistical significance between conditions was calculated by the non-parametric ANOVA, Kruskall-Wallis with uncorrected Dunn’s test (ns = Not-Significant, P > 0.05, * P < 0.05, ** P < 0.01, *** P < 0.001).

The lungs of YF-S0 vaccinated animals remained normal, or near to normal with no more signs of bronchopneumonia which is markedly different to sham-vaccinated animals (n=12 from two independent experiments; Figure 4). The specific disease scores and biomarkers quantified^2^ included (*i*) a reduction or lack of detectable lung pathology as observed by histological inspection (Figure 4A,B, Figure S6A); and, (*ii*) a significant improvement of the individual lung scores (p = 0.002) (Figure 4C, Figure S6B) and respiratory capacity (*i*.*e\** 32% less of lung volume obstructed; p = 0.0323; Figure 4D) in YF-S0 vaccinated animals as derived by micro-computed tomography (micro-CT) of the chest. In addition, immunization with YF-S0 resulted in an almost complete, in most cases full, normalization of the expression of cytokines, *e*.*g*., IL-6, IL-10, or IFN-γ in the lung, linked to disease exacerbation in COVID-19 (Figure 4E,F and Figure S7)^19-21^. Even the most sensitive markers of viral infection, such as the induction of antiviral Type III interferons (IFN-λ)^22^, or the expression of IFN-stimulated genes (ISG) such as MX2 and IP-10 in YF-S0 vaccinated animals showed no elevation as compared to levels in the lungs of untreated healthy controls (Figure 4F and Figure S7).

Overall, YF-S0 that expresses the non-cleavable S variant outcompeted construct YF-S1/2 expressing the cleavable version of S. This argues for the stabilized prefusion form of S serving as a relevant protective antigen for SARS-CoV-2. Moreover, in line with its failure to induce nAbs (Figure 3B), construct YF-S1 expressing solely the hACE2 receptor-binding S1 domain (Figure 1D) did not confer any protection against SARS-CoV-2 challenge in hamsters (Figure 3E, F and Figure 4).

### Immunogenicity, in particular a favorable Th1 polarization of cell-mediated immunity in mice

Since there are very few tools available to study CMI in hamsters, humoral and CMI responses elicited by the different YF-S constructs were studied in parallel in mice. Since YF17D does not readily replicate in wild-type mice^23,24^, *Ifnar*^-/-^ mice that are deficient in Type I interferon signaling and that are hence susceptible to vaccination with YF17D, were employed^10,24,25^.

Mice were vaccinated with 400 PFU (of either of the YF-S variants, YF17D or sham) at day 0 and were boosted with the same dose 7 days later (Figure 5A). At day 21 all YF-S1/2 and YF-S0 vaccinated mice (n>9 in three independent experiments) had seroconverted to high levels of S-specific IgG and nAbs with log_10_ GMT of 3.5 (95% CI, 3.1-3.9) for IgG and 2.2 (95% CI, 1.7-2.7) for nAbs in the case of YF-S1/2, or 4.0 (95% CI, 3.7-4.2) for IgG and 3.0 (95% CI, 2.8-3.1) for nAbs in the case of YF-S0 (Figure 5B,C). Importantly, seroconversion to S-specific IgG was detectable as early as 7 days after the first immunization (Figure 5D). Isotyping of IgG revealed an excess of IgG2b/c over IgG1 indicating a dominant pro-inflammatory and hence antiviral (Th1) polarization of the immune response (Figure 5E) which is considered important for vaccine-induced protection against SARS-CoV-2^26-28^. Alike in hamsters, YF-S1 failed to induce SARS-CoV-2 nAbs (Figure 5B,C). However, high levels of YF nAbs were conjointly induced by all constructs confirming a consistent immunization (Figure S8).

**Fig. 5.**
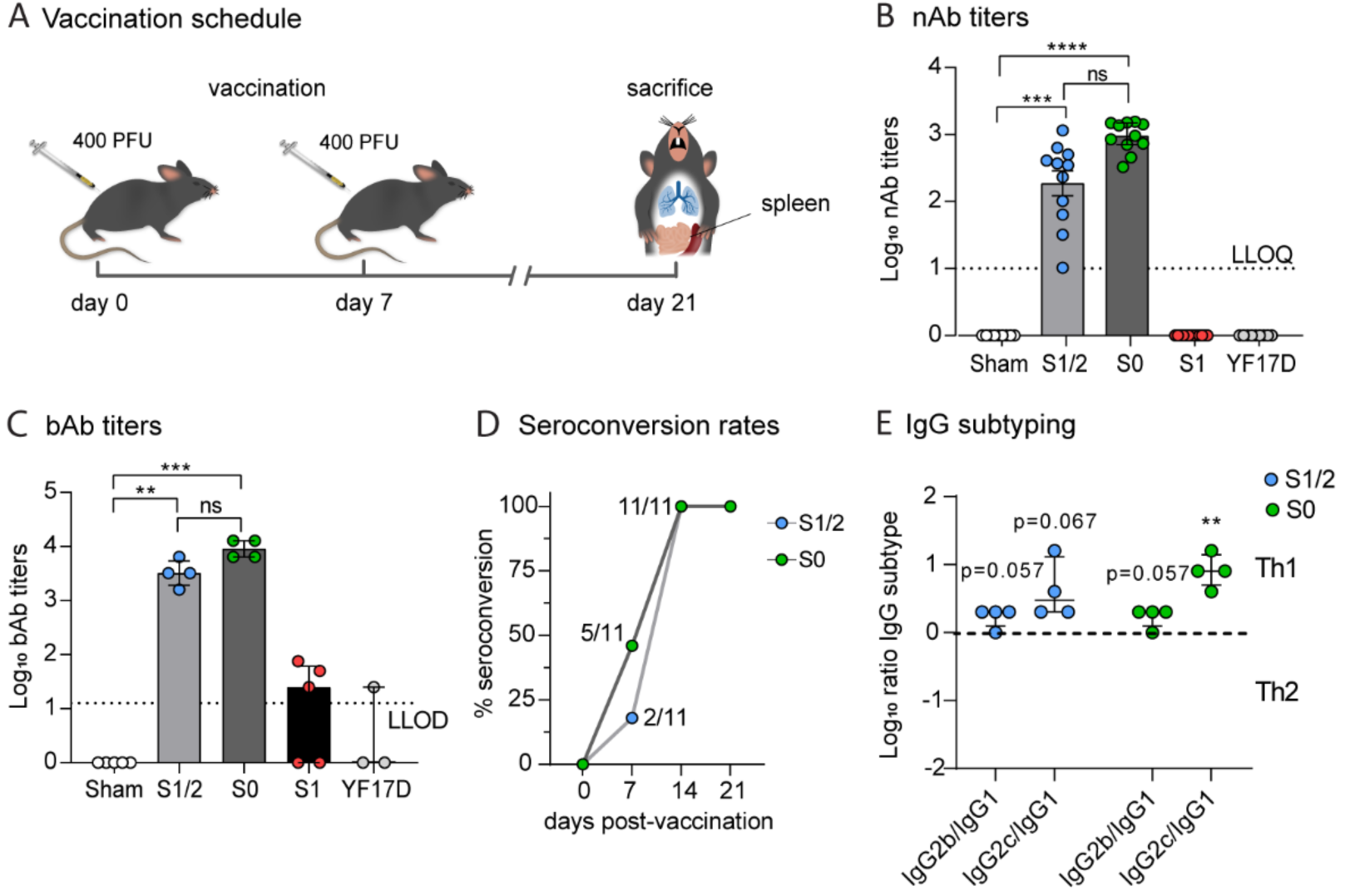
Humoral immune response elicited by YF-S vaccine candidates in mice. **(A) Schematic presentation of immunization and challenge schedule**. *Ifnar*^*-/-*^ mice were vaccinated twice i.p. with 400 PFU each at day 0 and 7 in five groups: constructs YF-S1/2 (blue, n=11), YF-S0 (green, n=11), YF-S1 (red, n=13), sham (white, n=9) or YF17D (grey, n=9). **(B, C) SARS-CoV-2 specific antibody levels**. Titers of nAbs **(B)** and bAbs **(C)** at day 21 post-vaccination. **(D) Seroconversion rates**. Rates at indicated days post-vaccination with YF-S1/2 and YF-S0 (number of animals with detectable bAbs at each time point are referenced). For quantification of bAbs, minipools of sera of two to three animals each were analyzed. Dotted lines indicate lower limit of quantification (LLOQ) or lower limit of detection (LLOD). **(E) IgG for YF-S1/2 (blue) and YF-S0 (green)**. Ratios of IgG2b or IgG2c over IgG1 (determined for minipools of two to three animals each) plotted and compared to a theoretical limit between Th1 and Th2 response (dotted line indicates IgG2b/c over IgG1 ratio of 1). Data shown are medians ± IQR from three independent vaccination experiments (n > 9 for each condition). Statistical significance between groups was calculated by a non-parametric ANOVA, Kruskall-Wallis with uncorrected Dunn’s test (B-C) or parametric One-Sample T-test (D) (ns = Not-Significant, P > 0.05, * P < 0.05, ** P < 0.01, *** P < 0.001, **** P < 0.0001).

To assess SARS-CoV-2-specific CMI responses that play a pivotal role for the shaping and longevity of vaccine-induced immunity as well as in the pathogenesis of COVID-19^29,30^, splenocytes from vaccinated mice were incubated with a tiled peptide library spanning the entire S protein as recall antigen. In general, vaccination with any of the YF-S variants resulted in marked S-specific T-cell responses with a favorable Th1-polarization as detected by IFN-γ ELISpot (Figure 6A), further supported by an upregulation of T-bet (*TBX21*), in particular in cells isolated from YF-S0 vaccinated mice (p = 0.0198, n = 5). This CMI profile was balanced by a concomitant elevation of GATA-3 levels (*GATA3*; driving Th2; p = 0.016), but no marked overexpression of RORγt (*RORC*; Th17) or FoxP3 (*FOXP3*; Treg) (Figure 6B). Intriguingly, in stark contrast to its failure to induce nAbs in mice (Figure 5A,C), or protection in hamsters (Figure 2 and 3), YF-S1 vaccinated animals had a greater number of S-specific splenocytes (p < 0.0001, n = 7) than those vaccinated with YF-S1/2 or YF-S0 (Figure 6A). Thus, even a vigorous CMI may not be sufficient for vaccine efficacy. A more in-depth profiling of the T-cell compartment by means of intracellular cytokine staining (ICS) and flow cytometry confirmed the presence of S-specific IFN-γ and TNF-α expressing CD8^+^ T-lymphocytes, and of IFN-γ expressing CD4^+^ (Figure 6E) and γ/δ T lymphocytes (Figure 6F), in particular in YF-S0 immunized animals. A specific and pronounced elevation of other markers such as IL-4 (Th2 polarization), IL-17A (Th17), or FoxP3 (regulatory T-cells) was not observed for YF-S1/2 or YF-S0. This phenotype is supported by t-SNE plot analysis of the respective T-cell populations in YF-S1/2 and YF-S0 vaccinated mice (Figure 6G and S8 tSNE) showing an increased percentage of IFN-γ expressing cells. It further revealed, firstly, a similar composition of either CD4^+^ cell sets, comprising an equally balanced mixture of Th1 (IFN-γ^+^ and/or TNF-α^+^) and Th2 (IL-4^+^) cells, and possibly a slight raise in Th17 cells in the case of the YF-S0 vaccinated animals. Likewise, secondly, for both constructs the CD8^+^ T-lymphocyte population was dominated by IFN-γ or TNF-α expressing cells, in line with the matched transcriptional profiles (Figure 6B). Of note, though similar in numbers, both vaccines YF-S0 and YF-S1/2 showed a distinguished (non-overlapping) profile regarding the respective CD8^+^ T lymphocyte populations expressing IFN-γ. In fact, YF-S0 tended to induce S-specific CD8^+^ T-cells with a stronger expression of IFN-γ (Figure 6G and S8). In summary, YF-S0 induces a vigorous and balanced CMI response in mice with a favorable Th1 polarization, dominated by SARS-CoV-2 specific CD8^+^ T-cells expressing high levels of IFN-γ when encountering the SARS-CoV-2 S antigen.

**Fig. 6.**
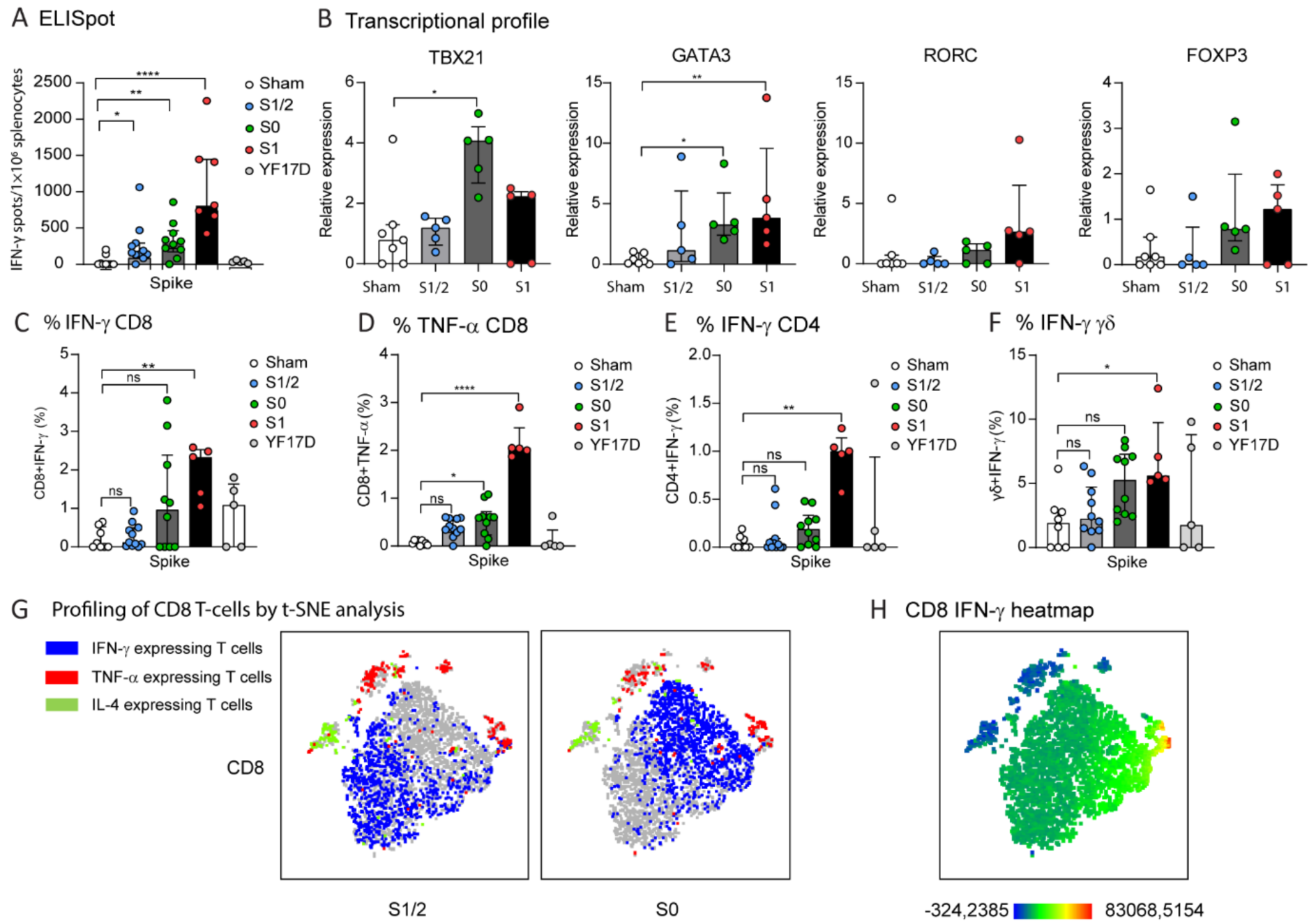
Cell-mediated immune (CMI) responses of YF-S vaccine candidates in mice. Spike-specific T-cell responses were analyzed by ELISpot and intracellular cytokine staining (ICS) of splenocytes isolated from *ifnar*^-/-^ mice 21 days post-prime (*i*.*e*., two weeks post-boost) immunization with YF-S1/2 (blue), YF-S0 (green), YF-S1 (red) in comparison to sham (white) or YF17D (grey). **(A) Quantitative assessment of SARS-CoV-2 specific CMI response by ELISpot**. Spot counts for IFN-γ-secreting cells per 10^6^ splenocytes after stimulation with SARS-CoV-2 Spike peptide pool. **(B) Transcriptional profile induced by YF-S vaccination**. mRNA expression levels of transcription factors (TBX21, GATA3, RORC, FOXP3) determined by RT-qPCR analysis of Spike peptide-stimulated splenocytes (n=5-7 per condition). Data were normalized for GAPDH mRNA levels and fold-changes over median of uninfected controls were calculated using the 2^(-ΔΔCq)^ method. **(C-F)** Percentage of IFN-γ **(C)** and TNF-α **(D)** expressing CD8, and IFN-γ expressing CD4 (**E**) and γ/δ **(F)** T-cells after stimulation with SARS-CoV-2 Spike peptide pool. All values normalized by subtracting spots/percentage of positive cells in corresponding unstimulated control samples. Data shown are medians ± IQR. Statistical significance between groups was calculated by the non-parametric ANOVA, Kruskall-Wallis with uncorrected Dunn’s test (ns = Not-Significant, P > 0.05, * P < 0.05, ** P < 0.01, *** P < 0.001). **(G, H) Profiling of CD8 T-cells from YF-S1/2 and YF-S0 vaccinated mice by t-SNE analysis**. t-distributed Stochastic Neighbor Embedding (t-SNE) analysis of spike-specific CD8 T-cells positive for at least one intracellular marker (IFN-γ, TNF-α, IL-4) from splenocytes of *ifnar*^-/-^ mice immunized with YF-S1/2 or YF-S0 (n=6 per group) after overnight stimulation with SARS-CoV-2 Spike peptide pool. Blue dots indicate IFN-γ expressing T-cells, red dots TNF-α expressing T-cells, and green dots IL-4 expressing CD8 T-cells. **(H) Heatmap** of IFN-γ expression density of spike-specific CD8 T-cells from YF-S1/2 and YF-S0 vaccinated mice. Scale bar represents IFN-γ expressing density (blue – low expression, red – high expression) (see Figure S8 for full analysis).

### Protection and short time to benefit after single-dose vaccination

Finally, vaccination of hamsters using a single-dose of YF-S0 induced high levels of nAbs and bAbs (Figure 7B and 7C) in a dose- and time-dependent manner. Furthermore, it appears a single 104 PFU dose of YF-S0 yielded higher levels of nAbs (log_10_ GMT 2.8; 95% CI: 2.5-3.2) at 21 days post-vaccination compared to the antibody levels in a prime-boost vaccination with two doses of 10^3^ PFU (log_10_ GMT 2.2; 95% CI: 1.9-2.6) (p = 0.039, two tailed Mann-Whitney test) (Figure 3B). Also, this single-dose regimen resulted in efficient and full protection against SARS-CoV-2 challenge, assessed by absence of infectious virus in the lungs in 8 out of 8 animals (Figure 7E). It should be noted that viral RNA at quantifiable levels was present in only 1 out of 8 animals (Figure 7D). In addition, protective immunity was mounted rapidly. Already 10 days after vaccination, 5 out of 8 animals receiving 10^4^ PFU of YF-S0 were protected against stringent infection challenge (Figure 7D and 7E).

**Fig. 7.**
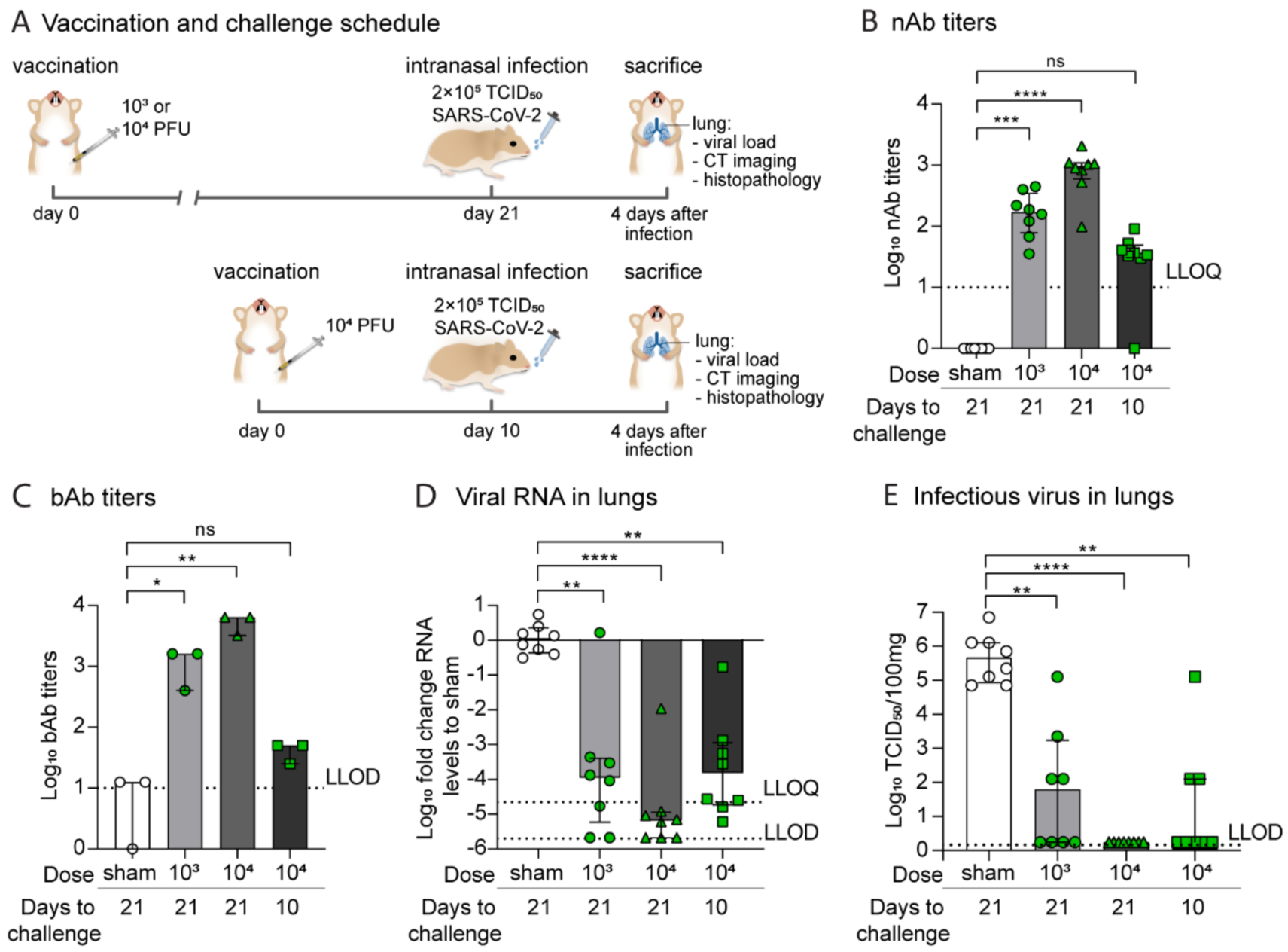
Single shot vaccination in hamsters using the YF-S0 lead vaccine candidate. **(A) Schematic presentation of experiment**. Three groups of hamsters were vaccinated only once i.p. with sham (white; n=8) or YF-S0 at two different doses; 1 × 10^3^ PFU (low, green circles; n=8) and 10^4^ PFU (high, green triangles; n=8) of YF-S0 at 21 days prior to challenge. A fourth group was vaccinated with the high 10^4^ PFU dose of YF-S0 at 10 days prior to challenge (green squares; n=8). **(B-C) Humoral immune responses following single dose vaccination**. Titers of nAb **(B)** and bAb **(C)** in sera collected from vaccinated hamsters immediately prior to challenge (minipools of sera of two to three animals analyzed for quantification of bAb). **(D, E) Protection from SARS-CoV-2 infection**. Protection from challenge with SARS-CoV-2 after vaccination with YF-S0 in comparison to sham vaccinated animals, as described for two-dose vaccination schedule (Figure 3 and Figure S5); log_10_-fold change relative to sham vaccinated in viral RNA levels **(D)** and infectious virus loads **(E)** in the lung of vaccinated hamsters at day four p.i. as determined by RT-qPCR and virus titration, respectively. Dotted line indicating lower limit of quantification (LLOQ) or lower limit of detection (LLOD) as indicated. Data shown are medians ± IQR. Statistical significance between groups was calculated by the non-parametric ANOVA, Kruskall-Wallis with uncorrected Dunn’s test (ns = Non-Significant, P > 0.05, * P < 0.05, ** P < 0.01, **** P < 0.0001).

## Discussion

Vaccines against SARS-CoV-2 need to be safe and result rapidly, ideally after one single dose, in long-lasting protective immunity. Different SARS-CoV-2 vaccine candidates are being developed, and several are vector-based. We report encouraging results of YF17D-vectored SARS-CoV-2 vaccine candidates. The post-fusion (S1/2), pre-fusion (S0) as well as the RBD S1 domain (S1) of the SARS-CoV-2 Spike protein were inserted in the YF17D backbone to yield the YF-S1/2, YF-S0 and YF-S1, respectively (Figure S1). The YF-S0 vaccine candidate, in particular, resulted in a robust humoral immune response in both, mice and Syrian hamsters.

Since SARS-CoV-2 replicates massively in the lungs of infected Syrian hamsters and results in major lung pathology^2,31-33^ we selected this model to assess the potency of these three vaccine candidates. YF-S0 resulted in efficient protection against stringent SARS-CoV-2 challenge, comparable, if not more vigorous, to other vaccine candidates in non-human primate models^16,17,34^. In about 40% of the YF-S0 vaccinated animals no increase in nAb levels (> 2x) following SARS-CoV-2 challenge was observed, suggestive for sterilizing immunity (no anamnestic response). In experiments in which animals were challenged three weeks after single 10^4^ PFU dose vaccination, no infectious virus was detected in the lungs. Considering the severity of the model, it is remarkable, that in several animals that were challenged with SARS-CoV-2 already 10 days after vaccination no infectious virus could be recovered from the lungs.

Reduction of viral replication mitigated lung pathology in infected animals with a concomitant normalization of biomarkers associated with infection and disease (Figure 4 and S6). Likewise, in lungs of vaccinated and subsequently challenged hamsters no elevation of cytokines, such as IL-6, was noted (Figure 4F).

Moreover, YF-S0 showed in two mice models a favorable safety profile as compared to the parental YF17D vector (Figure 2A and B). This is of importance as YF17D vaccine is contra-indicated in elderly and persons with underlying medical conditions. These preliminary, though encouraging, data suggest that YF-S0 might also be safe in those persons most vulnerable to COVID-19.

In addition, cell-mediated immunity (CMI) studied in mice revealed that YF-S0, besides efficiently inducing high titers of nAbs, favors a Th1 response. Such a Th1 polarization is considered relevant in light of a disease enhancement supposedly linked to a skewed Th2 immune^29^ or antibody-dependent enhancement (ADE)^35^. ADE may occur when virus-specific antibodies promote virus infection via various Fcγ receptor-mediated mechanisms, as suggested for an inactivated RSV post-fusion vaccine candidate^36^. A Th2 polarization may cause an induction and dysregulation of alternatively activated ’wound-healing’ monocytes/macrophages^26-28,37^ resulting in an overshooting inflammatory response (cytokine storm) thus leading to acute lung injury (ALI). No indication of such a disease enhancement was observed in our models.

In conclusion, YF-S0 confers vigorous protective immunity against SARS-CoV-2 infection. Remarkably, this immunity can be achieved within 10 days following a single dose vaccination. In light of the threat SARS-CoV-2 will remain endemic with spikes of re-infection, as a recurring plague, vaccines with this profile may be ideally suited for population-wide immunization programs.

## Methods

### Cells and viruses

BHK-21J (baby hamster kidney fibroblasts) cells^37^ were maintained in Minimum Essential Medium (Gibco), Vero E6 (African green monkey kidney, ATCC CRL-1586) and HEK-293T (human embryonic kidney cells) cells were maintained in Dulbecco’s Modified Eagle Medium (Gibco). All media were supplemented with 10% fetal bovine serum (Hyclone), 2 mM L-glutamine (Gibco), 1% sodium bicarbonate (Gibco). BSR-T7/5 (T7 RNA polymerase expressing BHK-21)^38^ cells were kept in DMEM supplemented with 0.5 mg/ml geneticin (Gibco).

For all challenge experiments in hamsters, SARS-CoV-2 strain BetaCov/Belgium/GHB-03021/2020 (EPI ISL 407976|2020-02-03) was used from passage P4 grown on Vero E6 cells as described^2^. YF17D (Stamaril^®^, Sanofi-Pasteur) was passaged twice in Vero E6 cells before use.

### Vaccine design and construction

Different vaccine constructs were generated using an infectious cDNA clone of YF17D (in an inducible BAC expression vector pShuttle-YF17D, patent number WO2014174078 A1)^10,12,39^. A panel of several SARS-CoV-2 vaccine candidates was engineered by inserting a codon optimized sequence of either the SARS-CoV-2 Spike protein (S) (GenBank: MN908947.3) or variants thereof into the full-length genome of YF17D (GenBank: X03700) as translational in-frame fusion within the YF-E/NS1 intergenic region^11,40^ (Figure S1). The variants generated contained (*i*) either the S protein sequence from amino acid (aa) 14-1273, expressing S in its post-fusion and/or prefusion conformation (YF-S1/2 and YF-S0, respectively), or (*ii*) its subunit-S1 (aa 14-722; YF-S1). To ensure a proper YF topology and correct expression of different S antigens in the YF backbone, transmembrane domains derived from WNV were inserted.

The SARS2-CoV-2 vaccine candidates were cloned by combining the S cDNA (obtained after PCR on overlapping synthetic cDNA fragments; IDT) by a NEB Builder Cloning kit (New England Biolabs) into the pShuttle-YF17D backbone. NEB Builder reaction mixtures were transformed into *E*.*coli* EPI300 cells (Lucigen) and successful integration of the S protein cDNA was confirmed by Sanger sequencing. Recombinant plasmids were purified by column chromatography (Nucleobond Maxi Kit, Machery-Nagel) after growth over night, followed by an additional amplification of the BAC vector for six hours by addition of 2 mM L-arabinose as described^10^.

Infectious vaccine viruses were generated from plasmid constructs by transfection into BHK-21J cells using standard protocols (TransIT-LT1, Mirus Bio). The supernatant was harvested four days post-transfection when most of the cells showed signs of CPE. Infectious virus titers (PFU/ml) were determined by a plaque assay on BHK-21J cells as previously described^10,14^. The presence of inserted sequences in generated vaccine virus stocks was confirmed by RNA extraction (Direct-zol RNA kit, Zymo Research) followed by RT-PCR (qScript XLT, Quanta) and Sanger sequencing, and by immunoblotting of freshly infected cells (see *infra*).

### Immunofluorescent staining

*In vitro* antigen expression of different vaccine candidates was verified by immunofluorescent staining as described previously by Kum et al. 2018. Briefly, BHK-21J cells were infected with 100 PFU of the different YF-S vaccine candidates. Infected cells were stained three days post-infection (3dpi). For detection of YF antigens polyclonal mouse anti-YF17D antiserum was used. For detection of SARS-CoV-2 Spike antigen rabbit SARS-CoV Spike S1 antibody (40150-RP01, Sino Biological) and rabbit SARS-CoV Spike primary antibody (40150-T62-COV2, Sino Biological) was used. Secondary antibodies were goat anti-mouse Alexa Fluor-594 and goat anti-rabbit Alexa Fluor-594 (Life Technologies). Cells were counterstained with DAPI (Sigma). All confocal fluorescent images were acquired using the same settings on a Leica TCS SP5 confocal microscope, employing a HCX PL APO 63x (NA 1.2) water immersion objective.

### Immunoblot analysis (Simple Western)

Infected BHK21-J cells were harvested and washed once with ice cold phosphate buffered saline, and lysed in radioimmunoprecipitation assay buffer (Thermo Fisher Scientific) containing 1x protease inhibitor and phosphatase inhibitor cocktail (Thermo Fisher Scientific). After centrifugation at 15,000 rpm at 4 °C for 10 minutes, protein concentrations in the cleared lysates were measured using BCA (Thermo Fisher Scientific). Immunoblot analysis was performed by a Simple Western size-based protein assay (Protein Simple) following manufactures instructions. Briefly, after loading of 400 ng of total protein onto each capillary, specific S protein levels were identified using specific primary antibodies (NB100-56578, Novus Biologicals and 40150-T62-CoV2, Sino Biological Inc.), and HRP conjugated secondary antibody (Protein Simple). Chemiluminescence signals were analyzed using Compass software (Protein Simple). To evaluate the removal of N-linked oligosaccharides from the glycoprotein, protein extracts were treated with PNGase F according to manufactures instructions (NEB).

### Animals

Wild-type Syrian hamsters (*Mesocricetus auratus*) and BALB/c mice and pups were purchased from Janvier Laboratories, Le Genest-Saint-Isle, France. *Ifnar1*^*-/-* 41^ and AG129^42^ were bred in-house. Six-to ten-weeks-old female mice and wild-type hamsters were used throughout the study.

### Animal Experiments

Animals were housed in couples (hamsters) or per five (mice) in individually ventilated isolator cages (IsoCage N – Biocontainment System, Tecniplast) with access to food and water *ad libitum*, and cage enrichment (cotton and cardboard play tunnels for mice, wood block for hamsters). Housing conditions and experimental procedures were approved by the Ethical Committee of KU Leuven (license P015-2020), following Institutional Guidelines approved by the Federation of European Laboratory Animal Science Associations (FELASA). Animals were euthanized by 100 µl (mice) or 500 µl (hamsters) of intraperitoneally administered Dolethal (200 mg/ml sodium pentobarbital, Vétoquinol SA).

### Immunization and infection of hamsters

Hamsters were intraperitoneally (i.p) vaccinated with the indicated amount of PFUs of the different vaccine constructs using a prime and boost regimen (at day 0 and 7). As a control, two groups were vaccinated at day 0 and day 7 with either 10^3^ PFU of YF17D or with MEM medium containing 2% FBS (sham). All animals were bled at day 21 to analyze serum for binding and neutralizing antibodies against SARS-CoV-2. At the indicated time after vaccination and prior to challenge, hamsters were anesthetized by intraperitoneal injection of a xylazine (16 mg/kg, XYL-M^®^, V.M.D.), ketamine (40 mg/kg, Nimatek^®^, EuroVet) and atropine (0.2 mg/kg, Sterop^®^) solution. Each animal was inoculated intranasally by gently adding 50 µl droplets of virus stock containing 2 × 10^5^ TCID_50_ of SARS-CoV-2 on both nostrils. Animals were monitored daily for signs of disease (lethargy, heavy breathing or ruffled fur). Four days after challenge, all animals were euthanized to collect end sera and lung tissue in RNA later, MEM or formalin for gene-expression profiling, virus titration or histopathological analysis, respectively.

### Immunization of mice

*Ifnar1*^*-/-*^ mice were i.p. vaccinated with different vaccine constructs by using a prime and boost of each 4 × 10^2^ PFU (at day 0 and 7). As a control, two groups were vaccinated (at day 0 and 7) with either YF17D or sham. All mice were bled weekly and serum was separated by centrifugation for indirect immunofluorescence assay (IIFA) and serum neutralization test (SNT). Three weeks post first-vaccination, mice were euthanized, spleens were harvested for ELISpot, transcription factor analysis by qPCR and intracellular cytokine staining (ICS).

### SARS-CoV-2 RT-qPCR

The presence of infectious SARS-CoV-2 particles in lung homogenates was quantified by qPCR^2^. Briefly, for quantification of viral RNA levels and gene expression after challenge, RNA was extracted from homogenized organs using the NucleoSpin™ Kit Plus (Macherey-Nagel), following the manufacturer’s instructions. Reactions were performed using the iTaq™ Universal Probes One-Step RT-qPCR kit (BioRad), with primers and probes (Integrated DNA Technologies) listed in Supplementary Table S1. The relative RNA fold change was calculated with the 2^-ΔΔCq^ method^43^using housekeeping gene β-actin for normalization.

### End-point virus titrations

To quantify infectious SARS-CoV-2 particles, endpoint titrations were performed on confluent Vero E6 cells in 96-well plates. Lung tissues were homogenized using bead disruption (Precellys^®^) in 250 µL minimal essential medium and centrifuged (10,000 rpm, 5 min, 4 °C) to pellet the cell debris. Viral titers were calculated by the Reed and Muench method^44^ and expressed as 50% tissue culture infectious dose (TCID_50_) per mg tissue.

### Histology

For histological examination, lung tissues were fixed overnight in 4% formaldehyde and embedded in paraffin. Tissue sections (5 μm) were stained with hematoxylin and eosin and analyzed blindly for lung damage by an expert pathologist.

### Micro-computed tomography (CT) and image analysis

To monitor the development of lung pathology after SARS-CoV-2 challenge, hamsters were imaged using an X-cube micro-computed tomography (CT) scanner (Molecubes) as described before^2^. Quantification of reconstructed micro-CT data were performed with DataViewer and CTan software (Bruker Belgium). A semi-quantitative scoring of micro-CT data was performed as primary outcome measure and imaging-derived biomarkers (non-aerated lung volume) as secondary measures, as previously described^2,45-48^.

### Neurovirulence in suckling mice and neurotropism in AG129 mice

BALB/c mice pups and AG129 mice were respectively intracranially or i.p. inoculated with the indicated PFU amount of YF17D and YF-S vaccine constructs and monitored daily for morbidity and mortality for 21 days post inoculation.

### Detection of total binding IgG and IgG isotyping by indirect immunofluorescent assay (IIFA)

To detect SARS-CoV-2 specific antibodies in hamster and mouse serum, an in-house developed indirect IFA (IIFA) was used. Using CRISPR/Cas9, a CMV-SARS-CoV-2-Spike-Flag-IRES-mCherry-P2A-BlastiR cassette was stably integrated into the ROSA26 safe harbor locus of HEK293T cells^49^. To determine SARS-CoV-2 Spike binding antibody end titers, 1/2 serial serum dilutions were made in 96-well plates on HEK293T-Spike stable cells and HEK293T wt cells in parallel. Goat-anti-mouse IgG Alexa Fluor 488 (A11001, Life Technologies), goat-anti-mouse IgG1, IgG2b and IgG2c Alexa Fluor 488 (respectively 115-545-205, 115-545-207 and 115-545-208 from Jackson ImmunoResearch) were used as secondary antibody. After counterstaining with DAPI, fluorescence in the blue channel (excitation at 386 nm) and the green channel (excitation at 485 nm) was measured with a Cell Insight CX5 High Content Screening platform (Thermo Fischer Scientific). Specific SARS2-CoV-2 Spike staining is characterized by cytoplasmic (ER) enrichment in the green channel. To quantify this specific SARS-CoV-2 Spike staining the difference in cytoplasmic vs. nuclear signal for the HEK293T wt conditions was subtracted from the difference in cytoplasmic vs. nuclear signal for the HEK293T SARS-CoV-2 Spike conditions. All positive values were considered as specific SARS-CoV-2 staining. The IIFA end titer of a sample is defined as the highest dilution that scored positive this way. Because of the limited volume of serum, IIFA end titers for all conditions were determined on minipools of two to three samples.

### Pseudotyped virus seroneutralization test (SNT)

SARS-CoV-2 VSV pseudotypes were generated as described previously^50-52^. Briefly, HEK-293T cells were transfected with a pCAGGS-SARS-CoV-2_Δ18_-Flag expression plasmid encoding SARS-CoV-2 Spike protein carrying a C-terminal 18 amino acids deletion^53,54^. One day post-transfection, cells were infected with VSVΔG expressing a GFP (green fluorescent protein) reporter gene (MOI 2) for 2h. The medium was changed with medium containing anti-VSV-G antibody (I1, mouse hybridoma supernatant from CRL-2700; ATCC) to neutralize any residual VSV-G virus input^55^. 24h later supernatant containing SARS-CoV-2 VSV pseudotypes was harvested.

To quantify SARS-CoV-2 nAbs, serial dilutions of serum samples were incubated for 1 hour at 37 °C with an equal volume of SARS-CoV-2 pseudotyped VSV particles and inoculated on Vero E6 cells for 18 hours. Neutralizing titers (SNT_50_) for YFV were determined with an in-house developed fluorescence based assay using a mCherry tagged variant of YF17D virus^10,39^. To that end, serum dilutions were incubated in 96-well plates with the YF17D-mCherry virus for 1h at 37 °C after which serum-virus complexes were transferred for 72 h to BHK-21J cells. The percentage of GFP or mCherry expressing cells was quantified on a Cell Insight CX5/7 High Content Screening platform (Thermo Fischer Scientific) and neutralization IC_50_ values were determined by fitting the serum neutralization dilution curve that is normalized to a virus (100%) and cell control (0%) in Graphpad Prism (GraphPad Software, Inc.).

### SARS-CoV-2 plaque reduction neutralization test (PRNT)

Sera were serially diluted with an equal volume of 70 PFU of SARS-CoV-2 before incubation at 37 °C for 1h. Serum-virus complexes were added to Vero E6 cell monolayers in 24-well plates and incubated at 37 °C for 1h. Three days later, overlays were removed and stained with 0.5% crystal violet after fixation with 3.7% PFA. Neutralization titers (PRNT_50_) of the test serum samples were defined as the reciprocal of the highest test serum dilution resulting in a plaque reduction of at least 50%.

### Antigens for T cell assays

PepMix™ Yellow Fever (NS4B) (JPT-PM-YF-NS4B) and subpool-1 (158 overlapping 15-mers) of PepMix™ SARS-CoV-2 spike (JPT-PM-WCPV-S-2) were used as recall antigens for ELISpot and ICS. Diluted Vero E6 cell lysate (50 μg/mL) and a combination of PMA (50 ng/mL) (Sigma-Aldrich) and Ionomycin (250 ng/mL) (Sigma-Aldrich) served as negative and positive control, respectively.

### Intracellular cytokine staining (ICS) and flow cytometry

Fresh mouse splenocytes were incubated with 1.6 μg/mL Yellow Fever NS4B peptide; 1.6 μg/mL Spike peptide subpool-1; PMA (50 ng/mL)/Ionomycin (250 ng/mL) or 50 μg/mL Vero E6 cell for 18h at 37 °C. After treatment with brefeldin A (Biolegend) for 4h, the splenocytes were stained for viability with Zombie Aqua™ Fixable Viability Kit (Biolegend) and Fc-receptors were blocked by the mouse FcR Blocking Reagent (Miltenyi Biotec)(0.5μL/well) for 15 min in the dark at RT. Cells were then stained with extracellular markers BUV395 anti-CD3 (17A2) (BD), BV785 anti-CD4 (GK1.5) (Biolegend), APC/Cyanine7 anti-CD8 (53-6.7) (Biolegend) and PerCP/Cyanine5.5 anti-TCR γ/δ (GL3) (Biolegend) in Brilliant Stain Buffer (BD) before incubation on ice for 25 min. Cells were washed once with PBS and fixed/permeabilized for 30 min by using the FoxP3 transcription factor buffer kit (Thermo Fisher Scientific) according to the manufacturer’s protocol. Finally, cells were intracellularly stained with following antibodies: PE anti-IL-4 (11B11), APC anti-IFN-γ (XMG1.2), PE/Dazzle™ 594 anti-TNF-α (MP6-XT22), Alexa Fluor^®^ 488 anti-FOXP3 (MF-14), Brilliant Violet 421 anti-IL-17A (TC11-18H10.1) (all from Biolegend) and acquired on a BD LSRFortessa™ X-20 (BD). All measurements were calculated by subtracting from non-stimulated samples (incubated with non-infected Vero E6 cell lysates) from corresponding stimulated samples. The gating strategy employed for ICS analysis is depicted in Fig. S9. The strategy used for comparative expression profiling of vaccine-induced T-cell populations by t-distributed Stochastic Neighbor Embedding (t-SNE) analysis is outlined in Fig. S8. All flow cytometry data were analysed using FlowJo Version 10.6.2 (LLC)). t-SNE plot was generated in Flowjo after concatenating spike-specific CD4 and CD8 T cell separately based on gated splenocyte samples.

### ELISpot

ELISpot assays for the detection of IFN-γ-secreting mouse splenocytes were performed with mouse IFN-γ kit (ImmunoSpot^®^ MIFNG-1M/5, CTL Europe GmbH). IFN-γ spots were visualized by stepwise addition of a biotinylated detection antibody, a streptavidin-enzyme conjugate and the substrate. Spots were counted using an ImmunoSpot^®^ S6 Universal Reader (CTL Europe GmbH) and normalized by subtracting spots numbers from control samples (incubated with non-infected Vero E6 cell lysates) from the spot numbers of corresponding stimulated samples. Negative values were corrected to zero.

### qPCR for transcription factor profile

Spike peptide-stimulated splenocytes split were used for RNA extraction by using the sNucleoSpin™ Kit Plus kit (Macherey-Nagel). cDNA was generated by using a high-capacity cDNA Reverse Transcription Kit (Thermo Fisher Scientific). Real-time PCR was performed using the TaqMan gene expression assay (Applied Biosystems) on an ABI 7500 fast platform. Expression levels of TBX21, GATA3, RORC, FOXP3 (all from Integrated DNA Technologies) were normalized to the expression of GAPDH (IDT). Relative gene expression was assessed by using the 2^-ΔΔCq^ method.

### Statistical analysis

GraphPad Prism (GraphPad Software, Inc.) was used for all statistical evaluations. The number of animals and independent experiments that were performed is indicated in the figure legends. Statistical significance was determined using the non-parametric Mann-Whitney U-test and Kruskal-Wallis test if not otherwise stated. Values were considered significantly different at P values?of ≤0.05.

## Acknowledgements

We thank Sarah Debayeve, Elke Maas, Jasper Rymenants, Tina Van Buyten and Caroline Collard (Laboratory of Virology, Rega) as well as Kathleen Van den Eynde, Eef Allegaert, Sarah Cumps and Wilfried Versin (Histology) for their outstanding technical support. We thank Catherina Coun, Jasmine Paulissen, Céline Sablon and Nathalie Thys (TVPC-Rega) for their excellent technical assistance for generating all serology data. We thank Jens Wouters and Dr. Johan Nuyts (MoSAIC and Nuclear Medicine and Molecular Imaging, KU Leuven) for help with micro-CT image analysis and support with imaging file processing. We thank Nele Berghmans, Sofie Knoops, Thy Pham and Helena Crijns from the Rega Immunology group (KU Leuven) and Nicolas Ongenae from ViroVet NV (Leuven) for help with hamster husbandry and bleeding.

We thank Julie Vercruysse, Cédric Vansalen and Dr. Nesya Goris (ViroVet NV, Leuven, Belgium) for help with large scale plasmid production. We thank Prof. Jelle Matthijnssens, Dr. Ward Deboutte and Lila Close for next-generation sequencing analysis of vaccine virus stocks. We thank Prof. Dirk Daelemans for access and Winston Chiu (Caps-It, Rega) for technical assistance with the High Content Screening Platform. This list of selfless people providing generous help during these exceptional times may not be complete. The corresponding authors may like to excuse for any oversight and thus omission.

For sharing materials, reagents and protocols we thank Prof. Michael A. Whitt (University of Tennessee Health Science CenterMemphis, TN, USA) for providing plasmids to rescue VSV-dG-GFP pseudoviruses; Dr. Hannah Kleine-Weber, Dr. Markus Hoffmann and Prof. Stefan Pöhlmann (DPZ, Göttingen, Germany) for sharing L1-Hybridoma supernatants and protocols for the generation VSV-pseudovirions; Prof. Berend Jan Bosch and Dr. Wentao Li (University of Utrecht, The Netherlands) for sharing SARS-CoV-S expression plasmids; Prof. Ian Goodfellow (University of Cambridge, UK) for providing BSR-T7 cells; Els Brouwers (PharmAbs, KU Leuven) for assistance in culturing hybridoma cells; Prof. Peter Bredenbeek (LUMC, The Netherland) for providing BHK-21J and Vero E6 cells. *Ifnar1*^*-/-*^ mouse breeding couples were a generous gift of Dr. Claude Libert, (IRC/VIB, University of Ghent, Belgium).

## Funding

This project has received funding from the European Union’s Horizon 2020 research and innovation program under grant agreements No 101003627 (SCORE project) and No 733176 (RABYD-VAX consortium), funding from Bill and Melinda Gates Foundation under grant agreement INV-00636, and was supported by the Research Foundation Flanders (FWO) under the Excellence of Science (EOS) program (VirEOS project 30981113), the FWO Hercules Foundation (Caps-It infrastructure), and the KU Leuven Rega Foundation. This project received funding from the Research Foundation – Flanders (FWO) under Project No G0G4820N and the KU Leuven/UZ Leuven Covid-19 Fund under the COVAX-PREC project. J.M. and X.Z. were supported by grants from the China Scholarship Council (CSC). C.C. was supported by the FWO (FWO 1001719N). G.V.V. acknowledges grant support from KU Leuven Internal Funds (C24/17/061) and K.D. grant support from KU Leuven Internal Funds (C3/19/057 Lab of Excellence). G.O. is supported by funding from KU Leuven (C16/17/010) and from FWO-Vlaanderen. We appreciate the in-kind contribution of UCB Pharma, Brussels.

Finally, we wish to express our gratitude to everyone who has put their weight behind the research. Whether they donated money, organized fundraising campaigns or helped spread the word. Every form of support has made the difference. Thanks to the impulse we have received through the COVID-19 Fund, we were strengthened in our effort to develop a vaccine. We therefore wish to thank all donors and volunteers for their continued support. For their unstinting generosity and above all, for their hopeful optimism.

## Declaration of Interests

L.S.F., J.N., and K.D. are named as inventors on patent describing the invention and use of coronavirus vaccines. All other authors have nothing to disclose.

## Notes

### Competing Interest Statement

The authors have declared no competing interest.

