## Supplementary figures for "A single-dose live-attenuated YF17D-vectored SARS-CoV2 vaccine candidate"

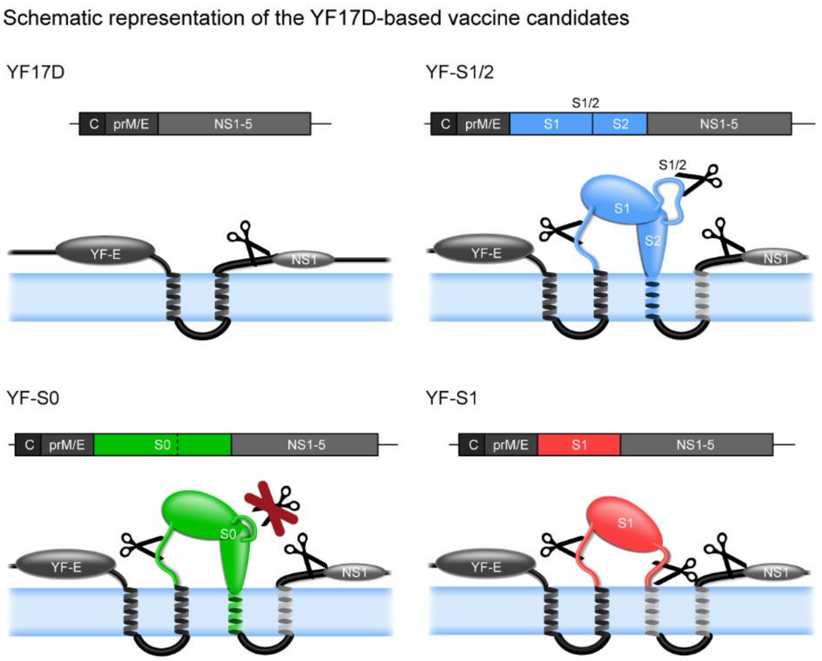


**Fig. S1. Schematic representation of the YF17D-based vaccine candidates (YF-S).** The SARS-CoV-2 Spike (S1/2, S0 or S1) antigen were inserted into the E/NS1 intergenic region as translational fusion within the YF17D polyprotein (dark grey) inserted in the ER (endoplasmic reticulum; pale blue). To cope with topological constraints of the fold of both SARS-CoV-2 Spike antigens and the polyprotein of the YF17D vector, one extra transmembrane domain (derived from the West Nile virus E-protein; light grey) was added to the C-terminal cytoplasmic domain of the full-length S proteins (S1/2 and S0). Likewise, two transmembrane domains were fused to the ER-resident C-terminus of the S1 subunit in construct YF-S1. Scissors indicate proposed maturation cleavage sites, including the S1/2 furin-cleavage site deleted in YF-S0.

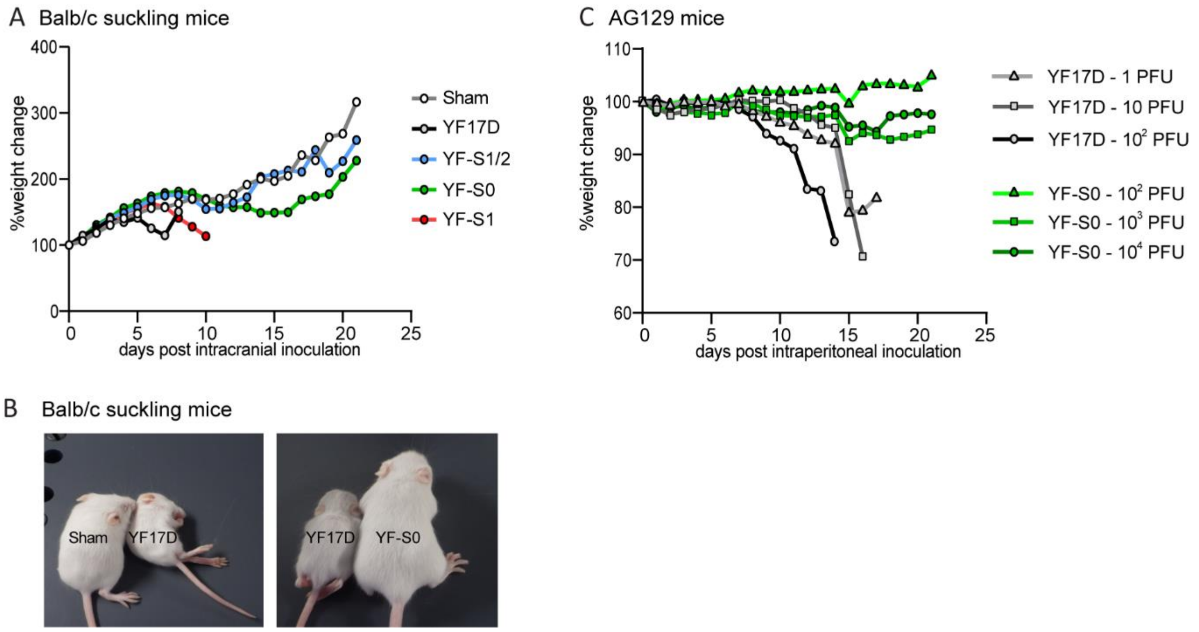


**Fig. S2. Attenuation of YF-S vaccine candidates. (A)** Weight evolution of suckling Balb/c mice (up to 21 days) after i.c. inoculation with 100 PFU of vaccine candidates (n=8) YF-S1/2 (blue), YF-S0 (green), YF-S1 (red) in comparison to sham (n=10, grey) or YF17D (n=9, black). **(B)** Representative images of Balb/c mice at seven days after intracranial inoculation with sham, $10^2$ PFU of either YF-S0 or YF17D. **(C)** Weight evolution of AG129 mice (up to 21 days) after intraperitoneal inoculation with a dose of $10^2$, $10^3$ or $10^4$ PFU of YF-S0 (green), and 1, 10 or $10^2$ PFU of YF17D (black and grey circles).

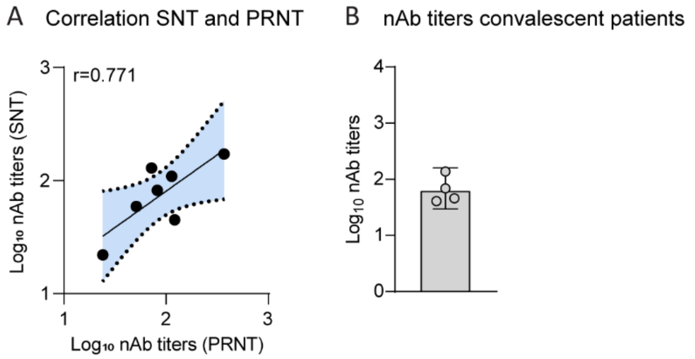


**Fig. S3. Correlation of nAb titers as determined by plaque reduction neutralization test (PRNT) and by serum neutralization test (SNT). (A)** Correlation analysis of nAb titers using SARS-CoV-2 (PRNT) and rVSV-ΔG-spike (SNT) for a panel of seven sera. $SNT_{50}$ and $PRNT_{50}$ values were plotted to determine the correlation between the neutralization assays with a Pearson regression coefficient of 0.77 (P=0.04). **(B)** NAbs in sera from four convalescent patients as determined by SNT. Data shown is median ± IQR.

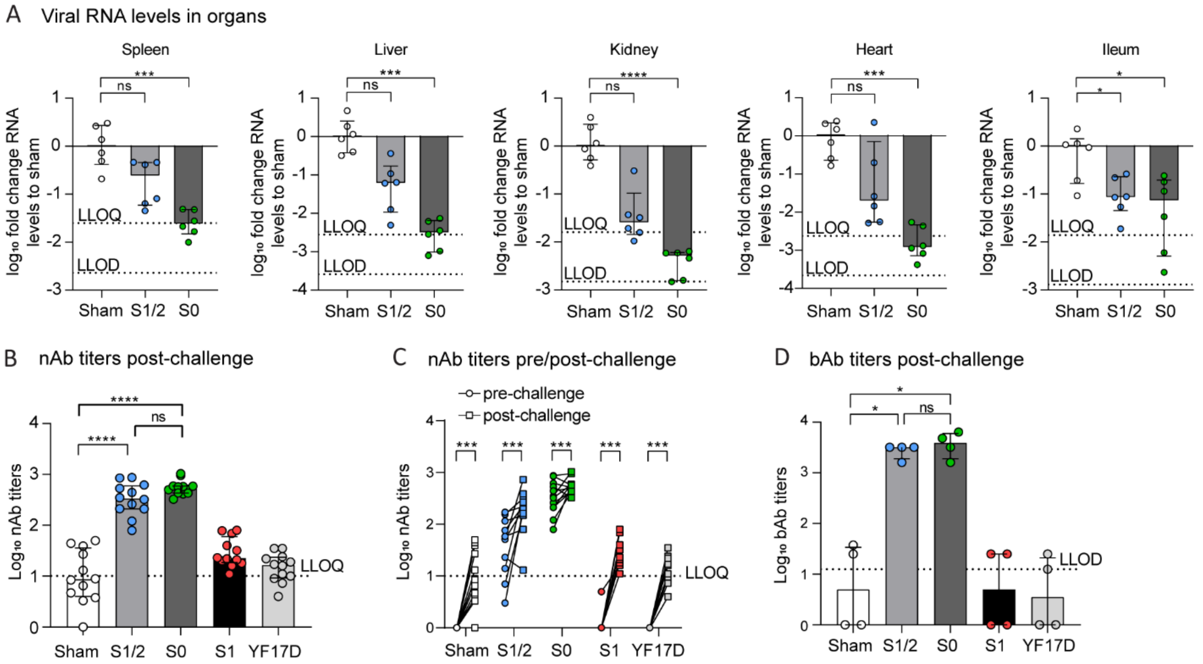


**Fig. S4. Immunogenicity and protective efficacy in hamsters. (A) Virus RNA load in organs.** Viral RNA in spleen, liver, kidney, heart and ileum of hamsters vaccinated with YF-S1/2, YF-S0 or sham, and challenged by infection with SARS-CoV-2. Viral RNA levels were determined by RT-qPCR, normalized against β-actin mRNA levels, and resulting fold-changes relative to the median of sham-vaccinated animals calculated using the $2^{(-\Delta\Delta Cq)}$ method. **(B-D) Anamnestic response.** NAb titers **(B)** and bAbs titers **(D)** in hamsters immunized with YF-S1/2 (green), YF-S0 (blue), YF-S1 (red) in comparison to sham (white) or YF17D (yellow) four days after challenge with SARS-CoV-2. **(C)** Pair-wise comparison of nAb titers of sera collected at day 21 post-immunization (circles), and four days post-challenge (squares). For quantification of bAbs, minipools of sera of three animals each were analyzed. Statistical significance between groups was calculated by the non-parametric ANOVA, Kruskall-Wallis with uncorrected Dunn's test (A, B and D), or a non-parametric Wilcoxon matched-pairs rank test (C) (ns = Not-Significant, P > 0.05, * P < 0.05, ** P < 0.01, *** P < 0.001, **** P < 0.0001).

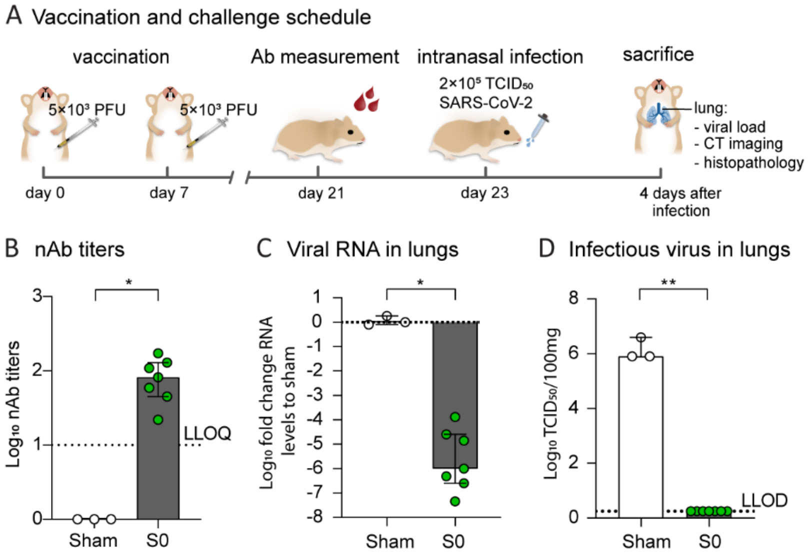


**Fig. S5. Immunogenicity and protective efficacy of vaccine candidate YF-S0 using a twice 5 x $10^3$ PFU dosing regimen. (A)** Schematic representation of immunization and challenge schedule. Syrian hamsters were immunized twice i.p. at day 0 and 7 with 5 x $10^3$ PFU each of vaccine constructs YF-S0 (green, n=7), sham (white, n=3). At day 23 post-vaccination, animals were intranasally inoculated with 2 × $10^5$ $TCID_{50}$ of SARS-CoV-2 and followed up for four days. **(B) Humoral immune responses.** NAb titers 21 days post-vaccination. **(C, D) Protection from SARS-CoV-2 infection.** Viral loads in lungs of hamsters four days after intranasal infection were quantified by RT-qPCR **(C)** and virus titration **(D)** as in Figure 3. Dotted line indicating lower limit of quantification (LLOQ) or lower limit of detection (LLOD) as indicated. Data shown are medians ± IQR. Statistical significance between groups was calculated by the non-parametric two-tailed Mann-Whitney test (* P < 0.05, ** P < 0.01).

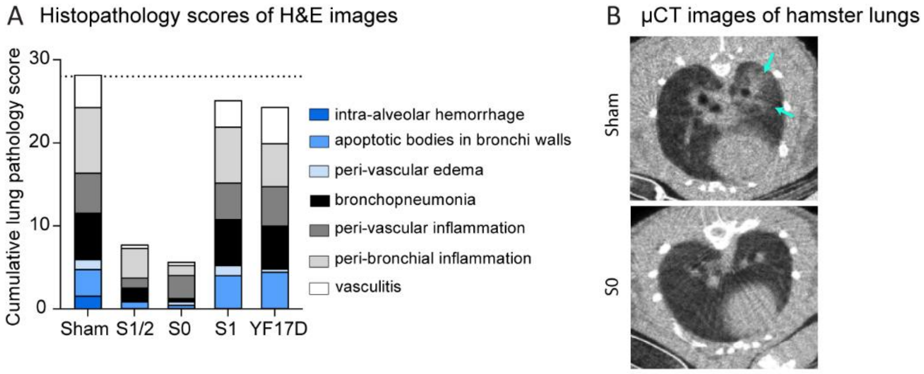


**Fig. S6. Lung pathology by histology and micro-CT imaging. (A)** Cumulative histopathology score for signs of lung damage (vasculitis, peri-bronchial inflammation, peri-vascular inflammation, bronchopneumonia, peri-vascular edema, apoptotic bodies in bronchus walls and intra-alveolar hemorrhage) in H&E stained lung sections (dotted line – maximum score in sham vaccinated group). **(B)** Representative micro-CT images of sham and YF-S0 vaccinated four days after SARS-CoV-2 infection. Arrows indicate examples of pulmonary infiltrates seen as consolidation of lung parenchyma (cyan).

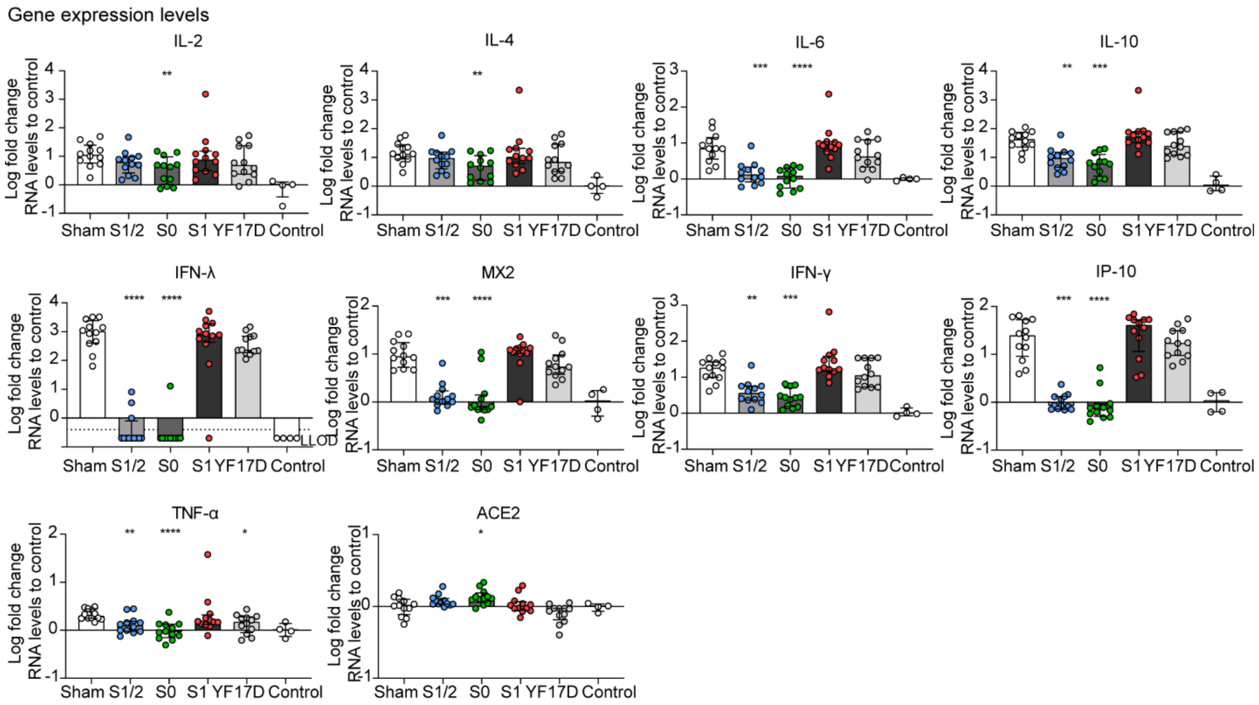


**Fig. S7. RNA expression levels after SARS-CoV-2 challenge.** Individual expression profiles for 10 genes in lungs of vaccinated hamsters (n=12 per group) four days after SARS-CoV-2 infection (as in Figure 4E) presented as $\log_{10}$-fold change relative to uninfected controls (n=4). Levels of individual mRNAs were determined by RT-qPCR and normalized for β-actin mRNA. Changes are reported as values over the median of uninfected controls calculated using the $2^{(-\Delta\Delta Cq)}$ method. Only for IFN-λ, where all control animals had undetectable RNA levels, fold changes were calculated over the lowest detectable value. Data presented as median ± IQR. LLOD – lower limit of detection (dotted line). Statistical significance compared to sham-vaccinated animals was calculated by a non-parametric ANOVA, Kruskall-Wallis with uncorrected Dunn's test (* $P < 0.05$, ** $P < 0.01$, *** $P < 0.001$, **** $P < 0.0001$).

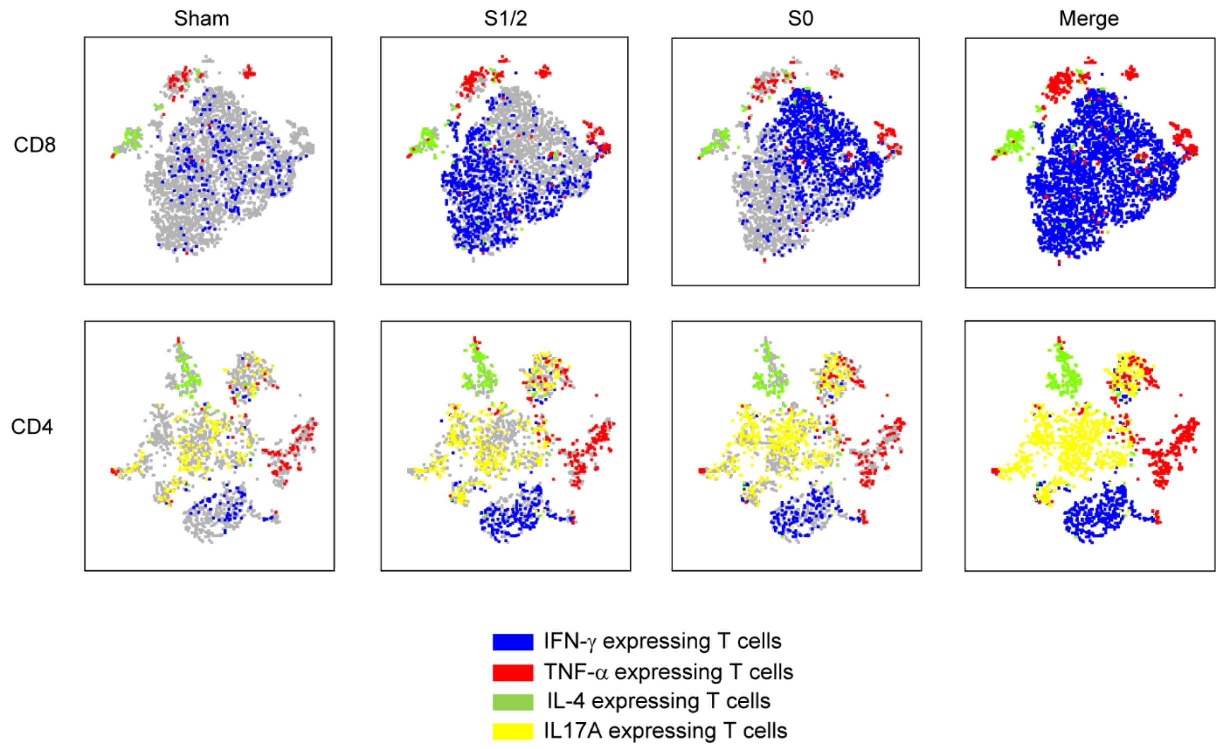


**Fig. S8. Profiling of CD8 and CD4 T-cells from YF-S1/2, YF-S0 and sham vaccinated mice by t-SNE analysis.** Full representation of t-distributed Stochastic Neighbor Embedding (t-SNE) analysis of Spike-specific CD4 and CD8 T-cells positive for at least one intracellular marker (IFN-γ, TNF-α, IL-4, or IL17A) from splenocytes of YF-S1/2, YF-S0 and sham vaccinated *ifnar*$^{-/-}$ mice (n=6 per group) after overnight stimulation with SARS-CoV-2 Spike peptide pool (red – IFN-γ expressing T-cells; blue – TNF-α expressing T-cells; green – IL-4 expressing T-cells; yellow – IL17A expressing T-cells). t-SNE plots generated using FlowJo by first concatenating Spike-specific CD8 (upper panels) or CD4 T-cells (lower panels) from all animals.

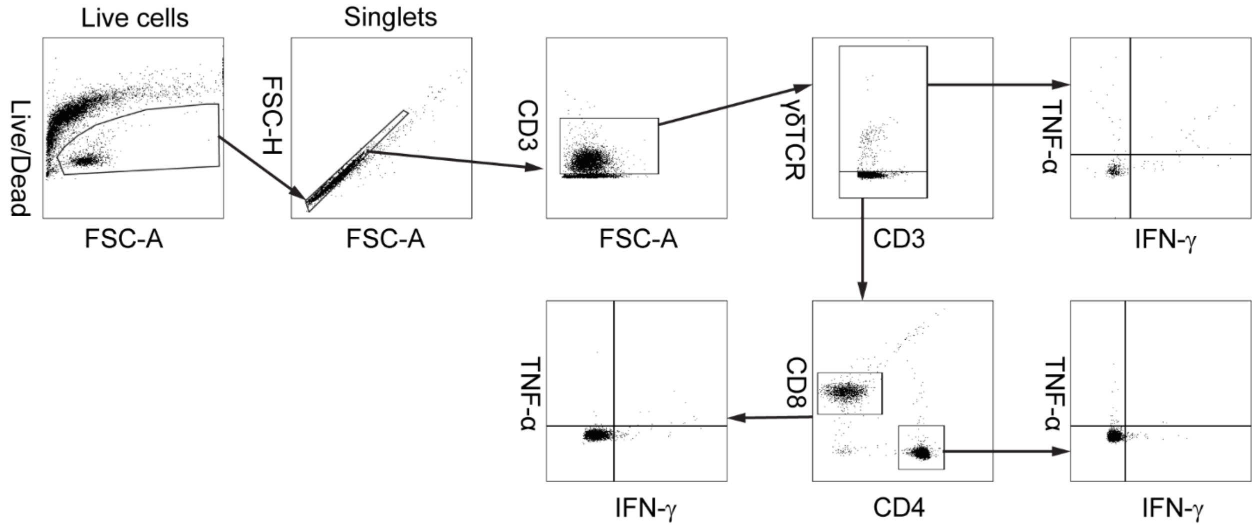


**Fig. S9. Sequential gating strategy for intracellular cytokine staining (ICS).** First, live cells were selected by gating out Zombie Aqua (ZA) positive and low forward scatter (FSC) events. Then, doublets were eliminated in a FSC-H *vs.* FSC-A plot. T-cells (CD3 positive) were stratified into γδT-cells (γδTCR⁺), CD4 T-cells (γδTCR⁻/CD4⁺) and CD8 T-cells (γδTCR⁻/CD8⁺). Boundaries defining positive and negative populations for intracellular markers were set based on non-stimulated control samples.

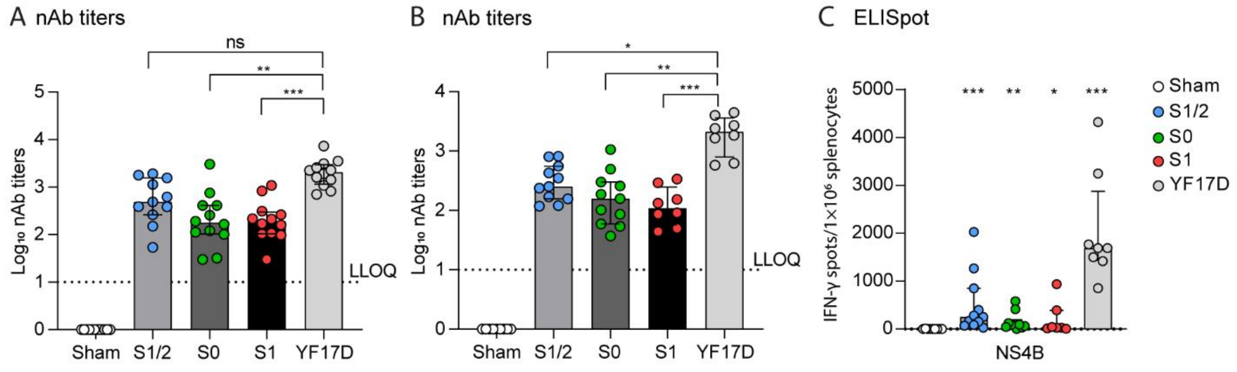


**Fig. S10. Humoral immune response elicited by YF in hamsters and mice. (A-B)** Neutralizing antibodies (nAb) in hamsters **(A)** and *ifnar$^{-/-}$* mice **(B)** vaccinated with the different vaccine candidates (sera collected at day 21 post-vaccination in both experiments (two-dose vaccination schedule). **(C) Quantitative assessment YF17D specific cell-mediated immune response by ELISpot.** Spot counts for IFNγ-secreting cells per $10^6$ splenocytes after stimulation with a NS4B peptide. Dotted line indicating lower limit of quantification (LLOQ) as indicated. Data shown are medians ± IQR. Statistical significance between groups was calculated by a non-parametric ANOVA, Kruskall-Wallis with uncorrected Dunn's test (ns = Not-Significant, P > 0.05, * P < 0.05, ** P < 0.01, *** P < 0.001).
