## Supplementary table 1 for "A single-dose live-attenuated YF17D-vectored SARS-CoV2 vaccine candidate"

**Supplementary Table S1**. Primers and probes used for RT-qPCR

| Gene | Description | Oligonucleotide sequence |
|---|---|---|
| **SARS-CoV-2** | Primer 1 | 5'-TTA CAA ACA TTG GCC GCA AA-3' |
| | Primer 2 | 5'-GCG CGA CAT TCC GAA GAA-3' |
| | Probe | 5'-FAM-ACA ATT TGC CCC CAG CGC TTC AG-BHQ1-3' |
| **Hamster *ACE2*** | Primer 1 | 5'-GGG AAC TGT CAA AGG GTA CAG-3' |
| | Primer 2 | 5'-CCC TTC CTA CAT CAG TCC TAC T-3' |
| | Probe | 5'-FAM-TCC CTG CTC ATT TGC TTG GTG ACA-ZEN/IABkFQ-3' |
| **Hamster *ACTB*** | Primer 1 | 5'-GGC CAG GTC ATC ACC ATT-3' |
| | Primer 2 | 5'-GAG TTG AAT GTA GTT TCG TGG ATG-3' |
| | Probe | 5'-Cy5-TTT CCA GCC TTC CTT CCT GGG TAT G-IBRQ-3' |
| **Hamster *IFN-γ*** | Primer 1 | 5'-TTT CTC CAT GCT GCT GTT GAA-3' |
| | Primer 2 | 5'-GGC CAT CCA GAG GAG CAT AG-3' |
| | Probe | 5'-FAM-CAC CAT CAA GGC AGA CCT GTT TGC TAA CTT-ZEN/IABkFQ-3' |
| **Hamster *IFNλ*** | Primer 1 | 5'-CCC ACC AGA TGC AAA GGA TT-3' |
| | Primer 2 | 5'-CTT GAG CAG CCA CTC TTC TAT G-3' |
| | Probe | 5'-FAM-ACA TAG CCC GGT TCA AGT CTC TGC-ZEN/IABkFQ-3' |
| **Hamster *IL-2*** | Primer 1 | 5'-AAG CTC CTG TAA GTC CAG CAG TAA C-3' |
| | Primer 2 | 5'-GTG CAC CCA CTT CAA GCT CTA A-3' |
| | Probe | 5'-FAM-AGG AAA CCC AGC AGC ACC TCG AGC-ZEN/IABkFQ-3' |
| **Hamster *IL-4*** | Primer 1 | 5'-GGG TCA CCT CAT GTT GGA AAT AAA-3' |
| | Primer 2 | 5'-CCA CGG AGA AAG ACC TCA TCT G-3' |
| | Probe | 5'-FAM-CAG GGC TTC CCA GGT GCT TCG CAA GT-ZEN/IABkFQ-3' |
| **Hamster *IL-6*** | Primer 1 | 5'-GGT ATG CTA AGG CAC AGC ACA CT-3' |
| | Primer 2 | 5'-CCT GAA AGC ACT TGA AGA ATT CC-3' |
| | Probe | 5'-FAM-AGA AGT CAC CAT GAG GTC TAC TCG GCA AAA-ZEN/IABkFQ-3' |
| **Hamster *IL-10*** | Primer 1 | 5'-TTC TGG CCC GTG GTT CTC T-3' |
| | Primer 2 | 5'-GTT GCC AAA CCT TAT CAG AAA TGA-3' |
| | Probe | 5'-FAM-CAG TTT TAC CTG GTA GAA GTG ATG CCC CAG G-ZEN/IABkFQ-3' |
| **Hamster *IP-10*** | Primer 1 | 5'-GCC ATT CAT CCA CAG TTG ACA-3' |
| | Primer 2 | 5'-CAT GGT GCT GAC AGT GGA GTC T-3' |
| | Probe | 5'-FAM-CGT CCC GAG CCA GCC AAC GA-ZEN/IABkFQ-3' |
| **Hamster *MX2*** | Primer 1 | 5'-CCA GTA ATG TGG ACA TTG CC-3' |
| | Primer 2 | 5'-CAT CAA CGA CCT TGT CTT CAG TA-3' |
| | Probe | 5'-FAM-TGT CCA CCA GAT CAG GCT TGG TCA-ZEN/IABkFQ-3' |
| **Hamster *TNF-$\alpha$*** | Primer 1 | 5'-AGC TGG TTG TCT TTG AGA GAC ATG-3' |
| | Primer 2 | 5'-GGA GTG GCT GAG CCA TCG T-3' |
| | Probe | 5'-FAM-CCA ATG CCC TCC TGG CCA ACG-ZEN/IABkFQ-3' |
| **Mouse *GAPDH*** | Primer 1 | 5'-GTG GAG TCA TAC GGA ACA TGT AG-3' |
| | Primer 2 | 5'-AAT GGT GAA GGT CGG TGT G-3' |
| | Probe | 5'-/56-FAM/TGC AAA TGG/ZEN/CAG CCC TGG TG/3IABkFQ/-3' |
| **Mouse *Tbx21*** | Primer 1 | 5'-CAA GAC CAC ATC CAC AAA CAT C-3' |
| | Primer 2 | 5'-TTC AAC CAG CAC CAG ACA G-3' |
| | Probe | 5'-/56-FAM/TCA CTA AGC/ZEN/AAG GAC GGC GAA TGT/3IABkFQ/-3' |
| **Mouse *GATA3*** | Primer 1 | 5'-GTC CCC ATT AGC GTT CCT C-3' |
| | Primer 2 | 5'-CCT TAT CAA GCC CAA GCG AA-3' |
| | Probe | 5'-/56-FAM/TGT CCC TGC/ZEN/TCT CCT TGC TGC/3IABkFQ/-3' |
| **Mouse *RORC*** | Primer 1 | 5'-GAG GTG CTG GAA GAT CTG C-3' |
| | Primer 2 | 5'-TCT GCA AGA CTC ATC GAC AAG-3' |
| | Probe | 5'-/56-FAM/CTA GCC AAG/ZEN/CTG CCA CCC AAA G/3IABkFQ/-3' |
| **Mouse *FOXP3*** | Primer 1 | 5'-CTG TCT TCC AAG TCT CGT CTG-3' |
| | Primer 2 | 5'-CTG GTC TCT GCA GGT TTA GTG-3' |
| | Probe | 5'-/56-FAM/CTG TGC CTG/ZEN/GTA TAT GCT CCC GG/3IABkFQ/-3' |
